# A Novel LysR-Type Global Regulator RpvA Controls Persister Formation and Virulence in *Staphylococcus aureus*

**DOI:** 10.1101/861500

**Authors:** Jian Han, Zhe Liu, Tao Xu, Wanliang Shi, Xiaogang Xu, Sen Wang, Lei Ji, Yuanyuan Xu, Qi Peng, Weiping Li, Ying Zhang

## Abstract

*Staphylococcus aureus* is the leading cause of wound and nosocomial infections. Persister formation and virulence factors play crucial roles during *S. aureus* infection. However, the mechanisms of persister formation and its relationship to virulence in *S. aureus* are poorly understood. In this study, we screened a transposon mutant library and identified a LysR-type global transcriptional regulator NWMN_0037, which we called RpvA, for regulator of persistence and virulence, whose mutation leads to higher susceptibility to antibiotics ampicillin and norfloxacin and various stresses including oxidative stress, heat, and starvation in late exponential and early stationary phase. Interestingly, the *rpvA* mutant was highly attenuated for virulence compared with the parent *S. aureus* Newman strain as shown by a much higher lethal dose, reduced ability to survive in macrophages and to form abscess in the mouse model. Transcriptional profiling and metabolomic analysis revealed that RpvA could repress multiple genes including *gapR*, *gapA*, *tpi*, *pgm*, *eno*, *glpD*, and *acs* expression and enhance production of numerous intermediate metabolites including dihydroxyacetone phosphate, 2-phosphoglycerate, acetyl-CoA, glycerol 3-phosphate, L-glutamate in the cells. The differentially expressed genes and altered production of metabolites are distributed in global metabolism including carbohydrate metabolism, amino acid metabolism, energy metabolism and metabolism of cofactors and vitamins. These metabolic adjustments could cause the cell to go into dormancy, thus promoting *S. aureus* to convert to persisters. In addition, RpvA could upregulate the expression of virulence genes including *hla*, *hlgA*, *hlgB*, *hlgC*, *lukF*, *lukS*, *lukD*, *sea* and *coa*, and carotenoid biosynthesis genes (*crtI*, *crtM*, *crtN*). Gel shift assay confirmed that RpvA could bind to the promoters of candidate target genes *hla*, *hlgB* and *crtM*, thus promoting *S. aureus* virulence. Because of the important functions of the RpvA, it may serve as an attractive target for developing new drugs and vaccines to more effectively control *S. aureus* infections.

## Introduction

Infections with *Staphylococcus aureus* constitute a major risk to human health, and are a leading cause of acute skin conditions, pneumonia, food poisoning, osteomyelitis, meningitis, arthritis, and toxic shock syndrome. In addition to acute infections, *S. aureus* can cause persistent infections such as blood stream infections, endocarditis, biofilms infections and pose significant challenges for treatment (1). The persistent infections that are refractory to treatment are thought to be in part due to the bacteria entering into a nongrowing antibiotic-tolerant state (2, 3).

The persister phenomenon was discovered 70 years ago when penicillin was found to fail to completely sterilize a *Staphylococcal* culture, and the small fraction of bacteria that survived the lethal penicillin treatment were called “persisters” (4). Persisters display phenotypic and non-heritable antibiotic tolerance or persistence due to dormant state in non-growing bacteria, and differs from genetic antibiotic resistance in growing bacteria that grow in the presence of antibiotics and have stable genetic basis. Cultures grown up from persisters remain susceptible to the same antibiotic as the parent culture from which the persisters are derived (4). Persisters are found in virtually all bacteria, and persister bacteria in some pathogenic bacteria such as *Mycobacterium tuberculosis*, *Escherichia coli*, *Pseudomonas aeruginosa*, *S. aureus*, *Borrelia burgdorferi*, *Brucella abortus etc.*, are known to play important roles in causing persistent infections and post-treatment relapse, and require lengthy treatment and pose significant challenges for effective treatment and control of these infections (5–7).

The mechanisms of bacterial persister formation are complex and are governed by redundant genes whose expression depends on particular conditions (8). Persisters have increased expression of several toxin/antitoxin (TA) modules (8–10). A toxin binds to the target and blocks the function, and the antibiotics can no longer corrupt its function, leading to tolerance (11). The toxin can be neutralized by antitoxin through inhibition of toxin translation or avid binding(12). The TA modules of *E. coli* include *hipBA*, *relBE*, *mazEF*, *dinJ-yafQ*, *mqsRA*, *ccdAB*, *tisB-*istR-1, *yefM*-*yoeB etc.* (13–20). The guanosine tetraphosphate (ppGpp) is essential for HipA7 mediated persistence (21). Other reported pathways include trans-translation (*ssrA* and *smpB*) (22), energy production (*sucA*, *fumC sucB* and *ubiF*) (23, 24), stringent response (21), SOS response/DNA repair (LexA) (25, 26), the phosphate and cellular metabolism PhoU mediated pathway (27) and efflux/transporters (28). Persister formation is associated with ATP depletion. A decrease in ATP levels leads to drug tolerance in *E. coli*, *P. aeruginosa*, and *S. aureus*. Environmental stimuli trigger a complex switch from poly(dC)/RmlB to P-poly(dC)/RmlB or RmlB in *S. aureus*, and this leads to decrease in intracellular ATP levels, forming dormant cells (29–31). In addition, biofilm formation is also an important mechanism for persistence to antibiotics (32).

Most peristers are dormant cells with slow metabolism and are in a non-growing physiological state where essential targets that antibiotics corrupt are thought to be inactive (33). Persisters require metabolic homeostasis in order to maintain culturability, sustain minimal adenylate charge, and repair damage and participate in the entry, maintenance, and exit from the persister phenotype (34). Inhibition of stationary phase respiration reduces persister formation in *E. coli* (35). Our knowledge on molecular mechanisms of persister formation was mainly derived from *E. coli* (9, 11, 15, 21, 27). In recent years, there have been some studies on mechanisms of *S. aureus* persister formation. ClpC was found to be a critical factor in staphylococcal energy metabolism, stress regulation, and persisters survival (36). Glycerol metabolism is important for *S. aureus* persister formation (37). Knocking out *sucA* and *fumC* caused high level of persisters formation which indicates that energy-generating components could serve as a mechanism of *S. aureus* persister formation (24). Oxidative stress defense mechanism regulated by *msaABCR* operon is required for persistent and recurrent *S. aureus* infections (38). Stationary phase *S. aureus* persister formation is associated with low membrane potential (39), while phenol-soluble modulins limit *S. aureus* prsister cell formation (40). Deletion of *ctaB* can attenuate *S. aureus* growth and virulence in mice but enhanced the formation of persister cells in stationary phase (41).

Despite several studies have been carried out on the mechanisms of persister formation in *S. aureus*, our understanding of the genes and pathways involved in the persister formation and the regulatory mechanisms remain incomplete. The mechanisms of persister formation can vary among bacterial species. For example, TA modules and the stringent response, which are known mechanisms of persister formation in *E. coli*, have recently been challenged to have a similar role in *S. aureus* (24). This further increases the need for studying the mechanisms of persister formation in *S. aureus*.

The ability of *S. aureus* to cause diseases depends on a diverse range of virulence factors including cell-surface associated molecules such as fibronectin-, fibrinogen-, and immunoglobulin-cell wall binding proteins and capsular polysaccharides and secreted exoproducts including pore-forming toxins (PFTs) (α bi-component leukocidins γ-hemolysin, Panton Valentine leukocidin (PVL), LukED, and LukGH/AB), enterotoxins, toxic shock syndrome toxin-1 (TSST-1), exfoliative toxins, phenol soluble modulins, and multiple secreted tissue-damaging exoenzymes including coagulase, lipases, and nucleases (1, 42–46). These virulence factors allow *S. aureus* to colonize, persist, and disseminate within the host, evade the immune system, and are responsible for the classical and potentially lethal signs of the infection, such as abscesses and sepsis (47). The exquisite and precise coordination of protein expression during different stages of infection depend on a network of regulatory genes (global regulatory network). A number of exoproducts are expressed in exponential phase, while in post exponential phase, the synthesis of exoproteins with enzymatic activity (e.g., hemolysins, toxins, proteases, and lipase) predominates and is controlled by global regulatory systems.

It has been hypothesized that persister genes may overlap with virulence genes (28, 48). Few studies explored the relationship between virulence and persister formation, and mostly it remains unclear (41). In the present study, we identified by mutant library screen a new transcription factor NWMN_0037 which we named RpvA based on its involvement in both persister formation and virulence in *S. aureus*. We performed transcriptional profiling and metabolomic analysis to further analyze the mechanisms by which RpvA regulates persister formation and virulence by detection of changes in gene expression and metabolites.

## Materials and methods

### Culture media, antibiotics, and chemicals

Tryptic soy broth (TSB) (BBL^TM^) and tryptic soy agar (TSA) (Difco^TM^) were obtained from Becton Dickinson (BD). Saline (0.9% NaCl) was used in the starvation experiment. Ampicillin, norfloxacin, tetracycline, vancomycin, erythromycin, rifampin, gentamicin and hydrogen peroxide (H_2_O_2_) were obtained from Sigma-Aldrich Chemical Co., and their stock solutions were freshly prepared, filter-sterilized and used at appropriate concentrations as indicated. BALB/c mice were purchased from Lanzhou Veterinary Institute (China).

### Bacterial strains and culture conditions

Bacterial strains and plasmids used in this study are listed in Table 1. All *S. aureus* strains except RN4220 (r-) were derivatives of Newman strain. All *S. aureus* strains were cultivated in TSA and TSB at 37 °C. For *S. aureus* carrying plasmid pT181, tetracycline was used at 10 μg/mL. As for the persister assay, antibiotics were used at the following concentrations, ampicillin, 10 μg/mL; norfloxacin, 20 μg/mL.

**Table 1.**
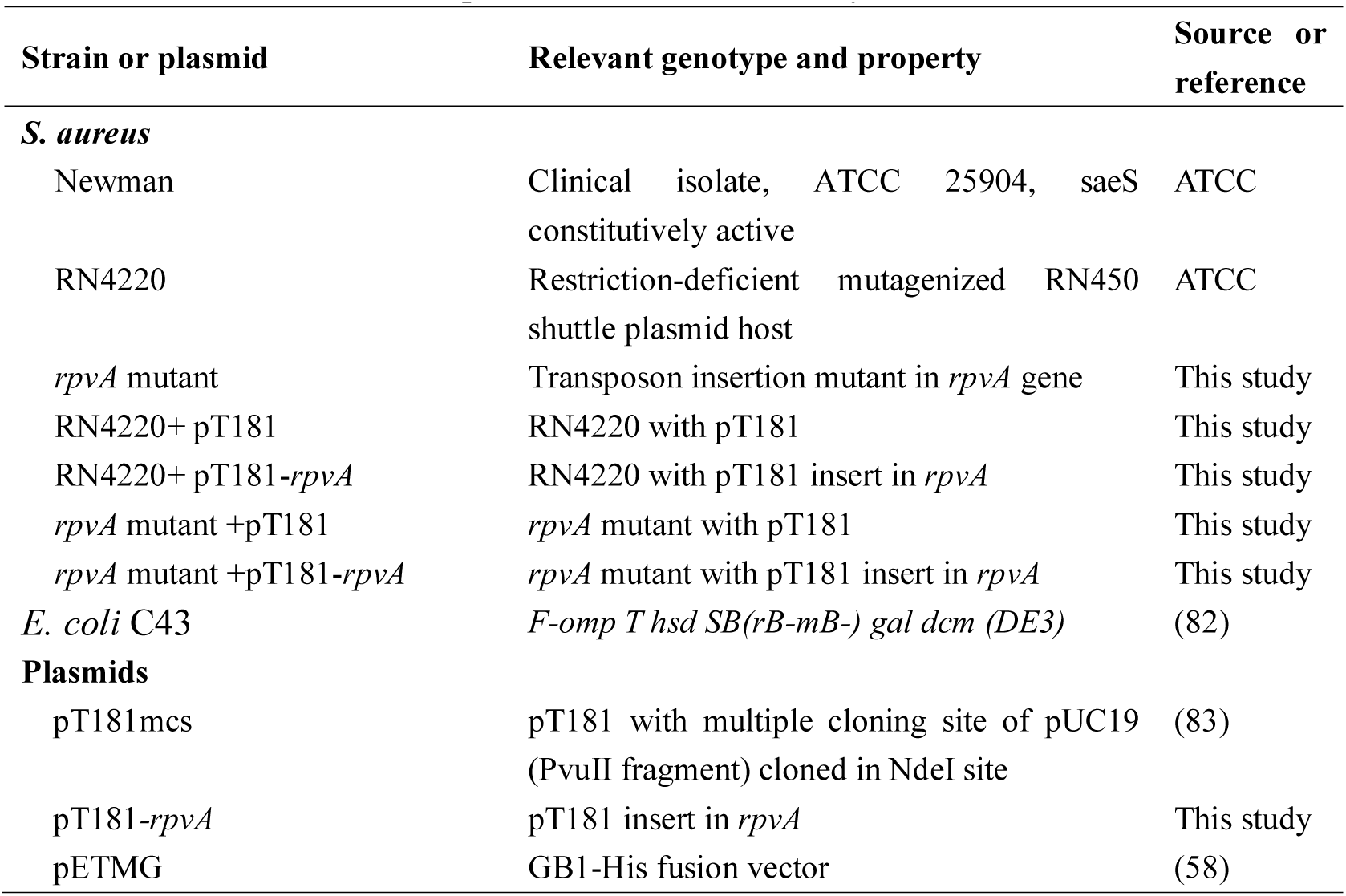
Bacterial strains and plasmids used in this study

### Genetic screens to identify mutants with defective persister survival

The *S. aureus* Newman transposon mutant library consisting of 6076 mutants which we had constructed previously (37) were thawed at room temperature and transferred to 96-well plate with 200 μL TSB by a 96-well replicator and grown at 37 °C overnight without shaking. The cultures were transferred to other two 96-well plates again with 200 μL TSB and incubated at 37 °C. Ampicillin at 10 μg/mL was added to one plate at 9 hours and another at 18 hours. After incubation for 3 and 6 days, the cultures were transferred to TSA agar plates by using a 96-well replicator and grown at 37 °C overnight. The mutants that did not grow at the above condition were picked from the corresponding original wells, cultured in TSB medium and rescreened by repeating the process above to confirm the phenotype.

### Inverse PCR and DNA sequencing

Genomic DNA was isolated from overnight cultures of persister deficient *S. aureus* mutants and the *S. aureus* Newman strain using lysostaphin (Sigma), glass beads (0.1mm), followed by phenol/chloroform extraction and ethanol DNA precipitation. The chromosomal DNA was digested by *Aci*I (New England Biolabs) and the DNA fragments were then circularized by using T4 DNA ligase (New England BioLabs). Primers ermF and ermR (Table 2) were used to amplify the circularized DNA in a 25 μL reaction volume and the PCR cycling parameters were 10 min at 96 °C, followed by 40 cycles of 30 s at 94 C, 30 s at 63 °C, and 3 min at 72 °C. The PCR products were sequenced with primer ermF and the DNA sequences were searched in the NCBI database using the BLAST algorithm to identify the gene of interest.

**Table 2.**
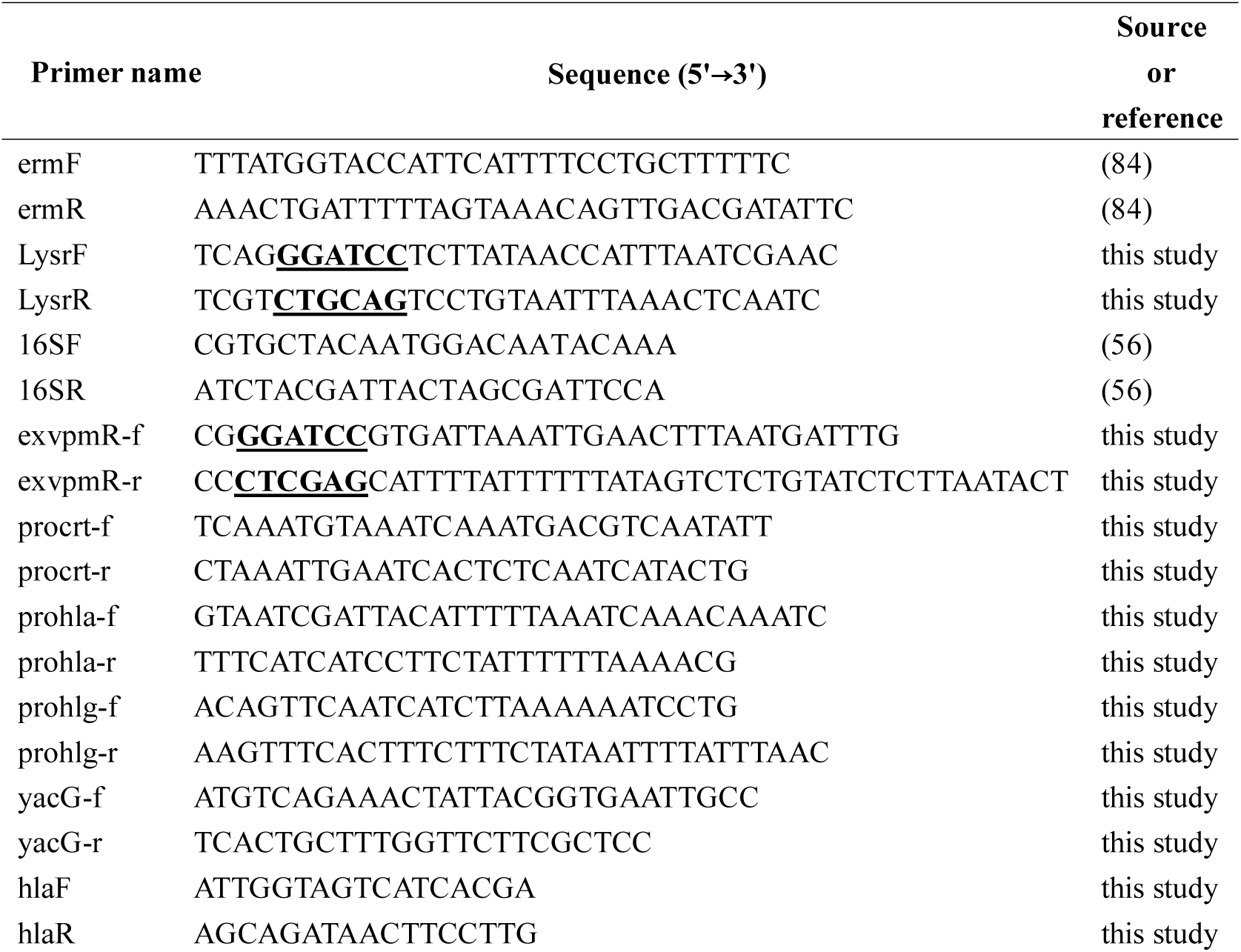
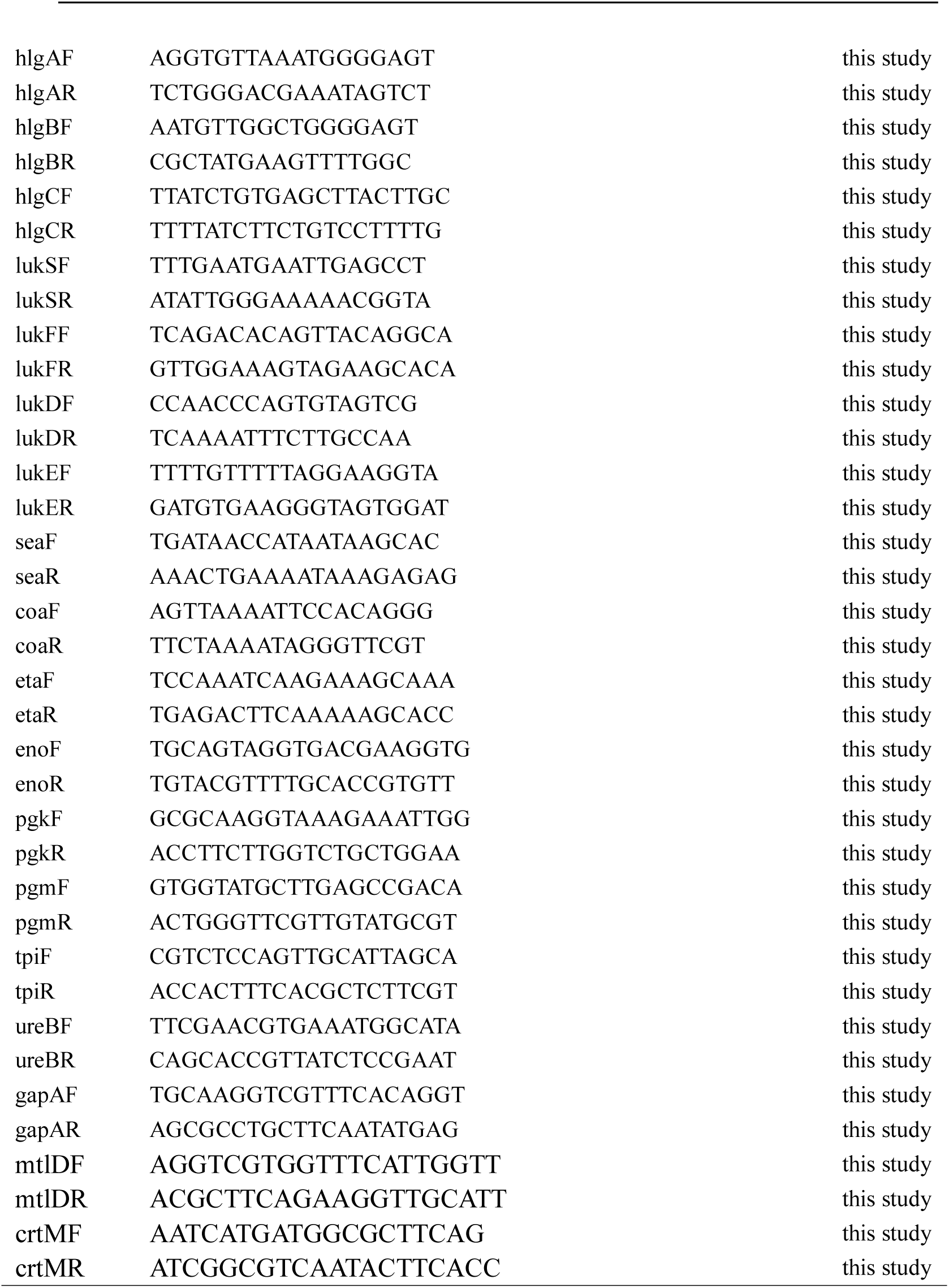
Oligonucleotides used in this study

### Complementation of mutant

For complementation of the *rpvA* mutant, the wild type *rpvA* gene from *S. aureus* Newman was amplified by PCR using primers LysrF and LysrR (Table2). The PCR primers were taken from −182 bp and 34 bp upstream and downstream of the *rpvA* gene and contained restriction sites *Bam*HI and *Pst*I, respectively. The PCR parameters were: 94 °C 15 min, followed by 30 cycles of 94 °C 30 s, 52 °C 30 s, and 72 °C 2 min, and a final extension at 72 °C for 10 min. After the PCR products were digested with *Bam*HI and *Pst*I, the DNA fragments were then cloned into plasmid pT181 cut with the same enzymes. The recombinant plasmid pT181-*rpvA* and the pT181 vector were electroporated into RN4220 (voltage = 2.5 kV, resistance = 100Ω, capacity = 25 μF) using MicroPulser^TM^ Electroporation Apparatus (Bio-Rad respectively. The transformed cells were spread on TSA plate (tet 5.0 μg/mL) incubated at 37 °C. The positive clones were identified by restriction digestion, and DNA sequencing. The plasmid pT181-r*pvA* and the pT181 vector control were then transformed into *S. aureus rpvA* mutant by electroporation as described.

### Minimum inhibitory concentration (MIC) and minimum bactericidal concentration (MBC) determination

MIC and MBC were determined using serial two-fold microdilution of the antibiotics (ampicillin, norfloxacin, rifampin, and vancomycin) in TSB liquid medium. The MIC was recorded as the minimum drug concentration that prevented visible growth of *S. aureus*. MBC was defined as 99.9% killing of the starting inoculum.

### Persister assays and stress exposure assays

The susceptibilities of the 3, 6, 9, 10, and 18-hour cultures of the wild type Newman strain, *rpvA* mutant + pT181 and the complemented strain to ampicillin (10μg/mL) or norfloxacin (20 μg/mL) were evaluated in drug exposure experiments in TSB. The cultures exposed to the above antibiotics were incubated without shaking at 37 °C for up to 1 week. Aliquots of cultures exposed to antibiotics were taken at different time points and serial 10-fold dilutions were plated on TSA and colony forming units (CFUs) were counted to determine survival of the bacteria after antibiotic exposure.

For heat stress, 9-hour early stationary phase cultures of Newman strain, *rpvA* mutant and its complemented strain were exposed to 58 °C in a water bath and the survival of bacteria was determined on TSA plates at 0, 30, 60, 90, and 120 min. For oxidative stress tests, 9-hour cultures were exposed to hydrogen peroxide (H_2_O_2_) at concentrations of 100 mM for 4 hours without shaking. The bacterial cells were washed with fresh TSB and the survival of bacteria was determined on TSA at 0, 2 and 4 hours. For carbon starvation, 48-hour stationary phase cultures in TSB were washed thrice with saline and resuspended to the same volume of saline. The cell suspensions were incubated at 37 °C for 2 weeks without shaking and the CFUs at different time points were determined.

### Intracellular anti-phagocytosis studies in THP-1 macrophages

THP-1 cells (ATCC TIB-202), a myelomonocytic cell line displaying macrophage-like activity (49), were stimulated with PMA (Phorbol 12 - myristate – 13 - acetate) at a final concentration of 100 ng/mL. Overnight cultures of the wild type Newman strain, *rpvA* mutant and the complemented strain were washed twice and resuspended in RPMI1640 (Life Technologies) and the bacteria diluted 1:1000 in 1 mL RPMI1640-10% fetal bovine serum (FBS) (Sigma) and adjusted to a concentration of 10^5-6^ CFU/mL. This suspension was then used to replace the culture medium of THP-1 cells to yield an initial macrophage-to-bacterium ratio of 0.25, and phagocytosis was allowed to occur at 37 °C for 1 hour. Triplicate wells were used for each time point. THP-1 cells were then washed thrice with prewarmed phosphate-buffered saline (PBS). RPMI1640-10% FBS with gentamicin (50 μg/mL) were added to the well and the plates were incubated at 37 °C, 5% CO_2_, and 95% humidity, to kill any extracellular bacteria. At times 0, 2, 4, 8, and 24 hours after bacterial addition, the macrophage cells were collected and lysed in 0.05% SDS, and serial 10-fold dilutions of macrophage cell lysates containing bacteria were plated on TSA to determine intracellular bacterial numbers.

### Median lethal dosage (LD50) determination (50, 51)

Four week old BALB/c mice with half of male and female were used for measuring LD_50_ of *S. aureus* wild type Newman strain, *rpvA* mutant and the complemented strain. Six groups of five mice per group were injected intraperitoneally with 0.6 mL bacterial suspension in PBS with two-fold decreasing bacterial doses (range 10^8^~10^9^ CFU/mL) from overnight stationary phase cultures. After 5-day observation, LD_50_ of each strain was calculated by the Reed-Muench method (52).

### Mouse abscess model and in vivo growth

Four week old BALB/c male mice were used for the subcutaneous abscess model as described (53, 54). Exponential phase cells of *S. aureus* wild type strain Newman, *rpvA* mutant and the complemented strain were prepared by diluting overnight cultures 1:100 into TSB, which were incubated at 37 °C with shaking until the optical density at 600 nm (OD600) reached 0.8. The bacterial cells were washed with PBS and diluted 1:20 in PBS and then mixed with an equal volume of autoclaved Cytodex-1 beads (131 to 220 μm; Sigma) in PBS. A 0.2 mL suspension of *S. aureus* cells were injected subcutaneously into each shaved flank of an anesthetized mouse. After 48 hours, the mice were euthanized, and the subcutaneous abscesses were removed aseptically and homogenized in 2 mL TSB. After 10-fold serial dilutions in TSB were prepared, the total numbers of bacteria recovered from the abscesses were determined by CFU counts on TSA plate.

### RNA-seq to detect differences in gene expression between wild type strain and rpvA mutant

Overnight stationary phase cultures of *S. aureus* Newman strain and the *rpvA* mutant were diluted (1:1000) by fresh TSB and incubated with shaking at 37 °C. Triplicate samples were harvested by centrifugation at 6-hour point and stocked at −80 °C. The final bacterial samples were submitted to Genergy Biotechnology (Shanghai) to extract total RNA for RNA sequencing. Total RNA was extracted by using Trizol, followed by chloroform extraction and isopropanol precipitation, dissolved in RNase-free water. rRNA was removed from the total RNA by using ribo-zero^TM^ magnetic kit (Epicentre^®^, USA). The rRNA-free RNA was used subsequently for RNA-seq library construction using the TruSeq® RNA LT Sample Prep Kit v2 (Illumina company, USA) followed with manufacturer’s protocol. For the cDNA purification and quantification step, the Ampure Beads (Beckman, USA) and Quant-iT™ PicoGreen^®^ dsDNA Assay Kit (Life company, USA) were used. Libraries were sequenced with 50-bp reads, using the standard HiSeq 2000 (Illumina) protocol. Illumina raw data provided by the GERALD (Illumina) software package were aligned to a FASTA file containing the *S. aureus* genome to change the data into Fastq Format. The results of RNA-seq were confirmed by qPCR.

### QPCR was used to verify the results of RNA-seq and the status of virulence gene expression

Total RNA from the wild type Newman strain and the *rpvA* mutant which cultured 6 hours in TSB were isolated according to the manufacturer’s instructions (Sangon Biotech). The primers for *eno*, *pgk*, *pgm*, *tpi*, *gapA*, *mtlD*, *crtM*, *hla*, and *hlgB* which were used to verify the results of RNA-seq and for *hla*, *hlgA*, *hlgB*, *hlgC*, *lukS*, *lukF*, *lukD*, *lukE*, *sea*, *coa*, and *eta* which were used to analyze the virulence factor expression were designed using Primer Express software (version 2.0; Applied Biosystems) (Table 2). The qPCR was performed in triplicate and the procedure referred to the method we have described previously (55). The expression of 16S rRNA was used as the control for estimating fold changes (56). In brief, total RNA was converted to cDNA using SuperScript III First-Strand synthesis (Takara Bio) and the real-time RT-PCR Cycling parameters were 95 °C for 30s, followed by 40 cycles of 5 s at 95 °C, 30 s at 60 °C. Threshold cycle (DDCt) were used to compare the relative expression levels of the interesting genes. The 2^−ΔΔ CT^ method (57) and Independent-Samples Student’s *t*-test was performed for analysis of relative gene expression data, and *P*-values < 0.05 were considered significant difference.

### Gel shift assay

Since attempt to purify soluble recombinant protein with the whole *rpvA* sequence from *E. coli* C43 (DE3) cultures with several recombinant plasmids was unsuccessful, the truncated *rpvA* segment (vpmR-s1) was amplified by PCR from *S. aureus* Newman chromosomal DNA using primers exvpmR-f and exvpmR-r (Table 2). The *vpmRs*1 fragment (1-81 aa) containing flanking restriction sites *Bam*HI and *Xho*I was cloned into pETMG (58), resulting in expression plasmid pETMG-VpmR-S1. A growing culture (at 37 °C) of C43 (DE3) with plasmid pETMG-VpmR-S1 was induced by adding 0.5 mM IPTG (isopropyl-1-thio-D-galactopyranoside) at an OD_600_ of 0.6. After 4 hours of additional incubation at 30 °C, the cells were harvested and digested with Bugbuster Master Mix (Merck Millipore). The mixture was then centrifuged at 15,000 g at 4 °C and the supernatant was filtered with a 0.22 μm filter (Merck Millipore). A Ni-NTA column was used to purify the recombinant GB1-VpmR-S1 on an AKTA system (GE). The flow through was analyzed on an SDS-PAGE gel after dialysis with Buffer A (20 mM Tris, 500 mM NaCl, pH 8.0). The concentration of GB1-VpmR-S1 was determined with Bradford assay (59). The gel shift assay was performed to determine the interaction of VpmR-S1 and the promoters of its putative target genes as described (60). According to the RNA-seq results, we chose 3 promoters for gel shift assay. While the virulence gene *hla* is single gene, *hlgB* lies in the downstream of *hlgC* in the “*hlg*” operon and the *NWMN_2465* is the first gene in the “*crt*” operon followed by *crtI*, *NWMN_2463*, *crtM* and *crtN* in this order. The promoters of *crt*, *hla* and *hlg* were amplified by PCR with primers procrt-f and procrt-r, prohla-f and prohla-r, prohlg-f and prohlg-r (Table 2) using *S. aureus* Newman chromosomal DNA as a template. A part of coding sequence of the *yacG* gene from *E. coli* W3110 strain was PCR amplified and included as a control. In each reaction 10 ng of promoter DNA was mixed with titrated GB1-VpmR-S1 protein (0, 0.1 and 0.2 μg) in a binding buffer (10 mM Tris, 1mM EDTA, 100mM KCl, 0.1mM DTT, 100 μg/mL BSA and 10% glycerol, pH 8 at 20 °C). The mixtures were incubated at 30 °C for 25 min and loaded onto a 10% polyacrylamide gel and run in Tris-acetate-EDTA electrophoresis buffer for 1.5 h. The gel was removed and stained with 0.05 Gel Red (Biotium, Inc.) in 0.1 M NaCl for 30 min. The band shifts were detected under a UV light and photographed.

### LC-MS/MS based metabolomic analysis

For sample collection and preparation, overnight cultures of the *rpvA* mutant and wild type Newman strain were diluted 1:1000 with fresh TSB and incubated at 37 °C with shaking at 200 rpm for 6-hours. Bacteria from 10 mL cultures were pelleted by centrifugation at 5000×g for 5 min. The supernatant was discarded and the pellet was washed three times with 1mL PBS. The final pellet was frozen by liquid nitrogen and stored at −80 °C for metabolite extraction. The treatments were repeated eight times for metabolomics analysis. The samples were processed as described (61). The samples were slowly thawed at 4 °C. 1 mL of pre-cooled methanol/acetonitrile/water solution (2:2:1, v/v) were added, followed by vortexing. Ultrasonic crushing was performed at a low temperature for 30 min, and then centrifuged at 14000 g, 4 °C for 20 min. The supernatant was dried in a vacuum centrifuge, and stored at −80 °C. For the UPLC-Q-TOF/MS analysis, the samples were re-dissolved in 100 μL acetonitrile/water (1:1, v/v) solvent. Quality control (QC) samples which were used to monitor the stability and reproducibility of the instrument analysis were prepared by pooling 10 μL of each sample.

For LC-MS/MS analysis, analyses were performed using an UHPLC (1290 Infinity LC, Agilent Technologies) coupled to a quadrupole time-of-flight (AB Sciex TripleTOF 6600) in Shanghai Applied Protein Technology Co., Ltd. A 2.1 mm × 100 mm ACQUIY UPLC BEH 1.7 µm column (waters, Ireland) were used for HILIC separation. The mobile phase contained 25 mM ammonium acetate and 25 mM ammonium hydroxide in water (A) and acetonitrile (B) were used in both electrospray ionization source (ESI) positive and negative modes. The gradient was 85% B for 1 min and was linearly reduced to 65% in 11 min, and then was reduced to 40% in 0.1 min and kept for 4 min, and then increased to 85% in 0.1 min, with a 5 min re-equilibration period employed. The gradient elution procedure was as follows: 0-1 min, 85% B; 1-12 min, B was linearly reduced to 65%; 12-12.1 min, B was linearly reduced to 40%; 12.1-15 min, B was maintained at 40%; 15-15.1 min, B was linearly varied to 85%; 15.1-20 min B was maintained at 85%. The auto sampler was maintained at 4 °C. The QC samples were inserted regularly to monitor the precision and stability of the method. The ESI source conditions were set as follows: ion source Gas1 as 60, ion source Gas2 as 60, curtain gas as 30, source temperature: 600c, IonSpray Voltage Floating (ISVF) ± 5500 V (both positive and negative modes). In MS only acquisition, the instrument was set to acquire over the m/z range 60-1000 Da, and the accumulation time for TOF MS scan was set at 0.20 s/spectra. TOF MS scan m/z range: 60-1000 Da, product ion scan m/z range: 25-1000 Da, TOF MS scan accumulation time 0.20 s/spectra, product ion scan accumulation time 0.05 s/spectra; The product ion scan is acquired using information dependent acquisition with high sensitivity mode selected; declustering potential: ±60 V (both positive and negative modes); collision energy: 35±15 eV; exclude isotopes within 4 Da, candidate ions to monitor per cycle: 6.

### Data processing and statistical data analysis

The raw MS data were converted to mzML files using ProteoWizard MSConvert, and processed using XCMS for peak alignment, retention time correction, and peak area extraction. Metabolite structure identification was achieved via accuracy mass (< 25 ppm) matching and secondary spectral matching. The results were compared with a laboratory standards database (Shanghai Applied Protein Technology Co., Ltd.).

Principal component analysis (PCA), partial least squares discriminant analysis (PLS-DA) and orthogonal partial least squares discriminant analysis (OPLS-DA) were carried out to determine the similarities of the samples between groups in SIMCA-P 14.1 (Umetrix, Umea, Sweden). The differential metabolites were screened based on the variable importance in the projection (VIP) value obtained from the OPLS-DA model and Student’s *t*-test on the raw data, and the metabolites with a VIP > 1.0 and *P*-value < 0.05 were considered statistically significant.

## Results

### Screening of the S. aureus mutant library and complementation of rpvA mutants

In an attempt to identify mutants with defect in persister formation, we screened the *S. aureus* transposon mutant library for deficiency in survival upon ampicillin (10 μg/mL) exposure. Five mutants were initially obtained after the mutant library screen. But after repeated screens, only the *rpvA* mutant consistently showed a defect in persister survival in 9-hour cultures and was therefore chosen for further analysis. Inverse PCR followed by DNA sequencing analysis indicated that the transposon was inserted at nucleotide position 442 of the *rpvA* gene which encodes a novel LysR family regulatory protein (Figure 1).

**Figure 1.**
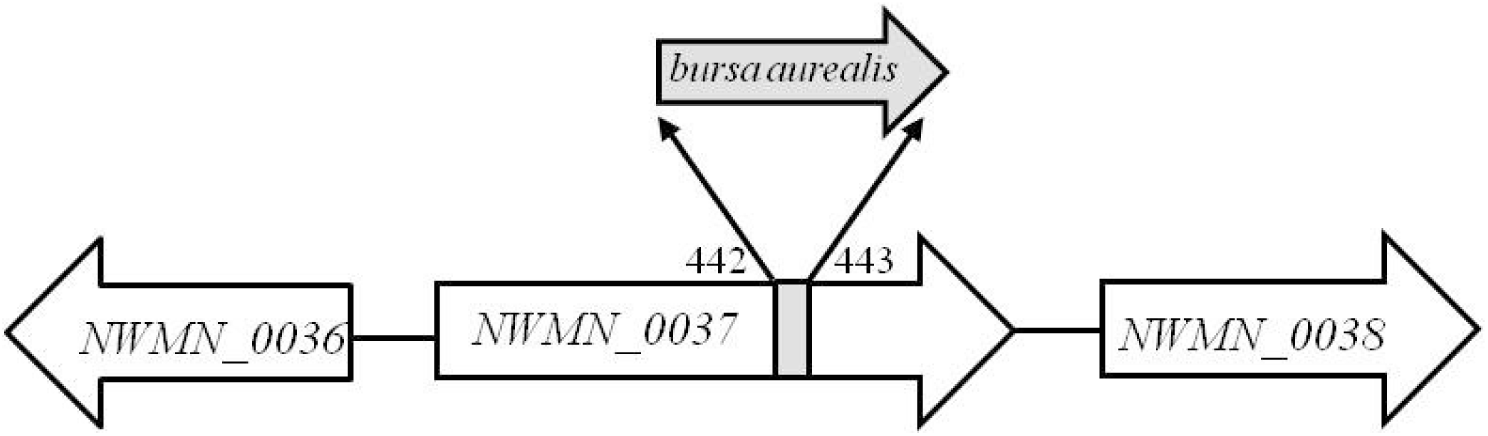
The region where the *bursa aurealis* transposon was inserted in *rpvA* (*NWMN_0037*) of *S. aureus* Newman DNA was between 442 and 443 bases.

Then recombinant plasmid pT181-*rpvA* was constructed and transformed into the *rpvA* mutant for complementation. The susceptibility to different antibiotics and stresses, and the virulence of the wild type, the *rpvA* mutant and its complemented strain were analyzed as below.

### Susceptibility of rpvA mutant and complemented strain to various antibiotics

To assess the susceptibility of the *rpvA* mutant to various antibiotics, including ampicillin, norfloxacin, rifampin, and vancomycin, both MIC and MBC experiments were performed with the wild type Newman strain as a control. The results showed that the *rpvA* mutant transformed with the vector was two-fold more susceptible to most antibiotics than the parent strain except vancomycin (Table 3). Complementation of the *rpvA* mutant with the wild type *rpvA* gene restored the wild-type level susceptibility to the antibiotics in both MIC and MBC tests (Table 3).

**Table 3.** MIC and MBC determination of the wild type, *rpvA* mutant and its complemented strain for different antibiotics.

| *S. aureus* | (MIC/MBC) ( $\mu\text{g mL}^{-1}$ ) | | | |
| --- | --- | --- | --- | --- |
|  | Ampicillin | Norfloxacin | Rifampin | Vancomycin |
| Newman | 0.315/0.315 | 2.0/2.0 | 0.02/0.02 | 4.0/4.0 |
| *rpvA* mutant+pT181 | 0.156/0.315 | 1.0/2.0 | 0.01/0.01 | 4.0/4.0 |
| *rpvA* mutant +pT181-*rpvA* | 0.625/0.625 | 1.0/2.0 | 0.02/0.02 | 4.0/8.0 |

To determine the effects of the *rpvA* mutation on bacterial survival upon drug exposure at different culture times (3, 6, 9, 10, and 18-hour), wild type Newman strain, the *rpvA* mutant and its complemented strain were exposed to ampicillin and norfloxacin, respectively. Antibiotic exposure assay showed that at early logarithmic phase (3 hours), the susceptibility of the *rpvA* mutant to ampicillin and norfloxacin was not significantly different from the wild type strain (Figure2A), but at late logarithmic phase stage (6 hours) and early stationary phase stage (9 hours), the *rpvA* mutant was more susceptible than the wild type. The *rpvA* mutant showed lower level of persisters than the wild type strain after 1 day or longer in ampicillin or norfloxacin exposure. Complementation of the *rpvA* mutant restored the level of persisters to that of the wild type (Figure 2 B and C). Interestingly, 9 hours was a critical time point, and the susceptibility of the *rpvA* mutant strain to antibiotics was significantly reduced from 10 hours, but there was no difference in susceptibility to antibiotics for the three strains when cultured for 18 hours or longer (Figure 2 D and E).

**Figure 2.**
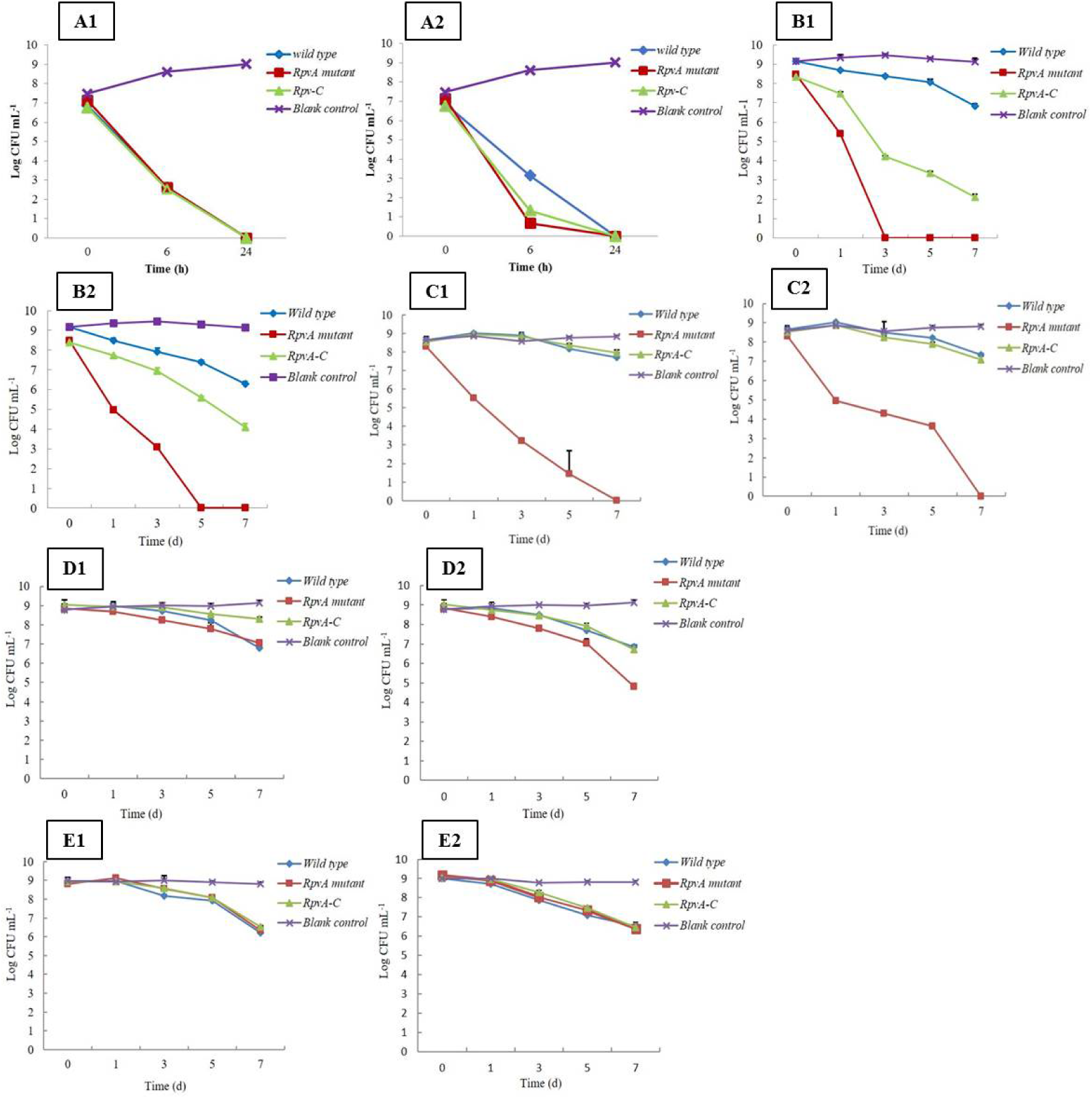
Susceptibilities of different growth phase cultures of *S. aureus* Newman wild type, the *rpvA* mutant, and the complemented strain upon ampicillin (10 μg/mL) and norfloxacin (20 μg/mL) exposure over time. A1, B1, C1, D1, and E1. 3, 6, 9, 10 and 18-hour cultures exposure to ampicillin. A2, B2, C2, D2, and E2. 3, 6, 9, 10 and 18-hour cultures exposure to norfloxacin.

### rpvA mutant was more susceptible to heat, starvation, and oxidative stress

To determine the effect of heat on the survival of the *rpvA* mutation, 9-hour cultures of the wild type, the *rpvA* mutant and its complemented strain were exposed to 58 ± 0.1°C. The *rpvA* mutant was much more sensitive to heat exposure than the others, and was completely killed after 2 hours of exposure, whereas the wild type and the complemented strain had 10^4^ and 10^3^ CFU mL^−1^ surviving bacteria, respectively (Figure 3 A). To determine the effect of hydrogen peroxide on the survival of the *rpvA* mutant, 9-hour cultures of the wild type, the *rpvA* mutant and its complemented strain were exposed to hydrogen peroxide (100 mM). Viable counts at different time points showed that the *rpvA* mutant was much more sensitive to H_2_O_2_ exposure than the wild type and the complemented strain, and was completely killed after 2 hours of exposure, whereas the wild type had 10^7^ CFU mL^−1^ surviving bacteria after 4 hours of exposure. The complemented strain restored peroxide resistance partially (Figure 3 B). To determine the effect of starvation on the survival of the *rpvA* mutation, 48-hour cultures of the wild type, the *rpvA* mutant and its complemented strain were suspended in saline and the survival of the bacteria was monitored at different time points. The results showed a more pronounced susceptibility to starvation of the *rpvA* mutant than the others. The survival of the *rpvA* mutant decreased 100-fold after 1 day of starvation, and no surviving bacteria were detected after 15 days, whereas the wild type strain still had 10^8^ and 10^6^ CFU mL^−1^ survivals after 1 day and 15 days of starvation. The complemented strain partially restored starvation resistance (Figure 3 C).

**Figure 3.**
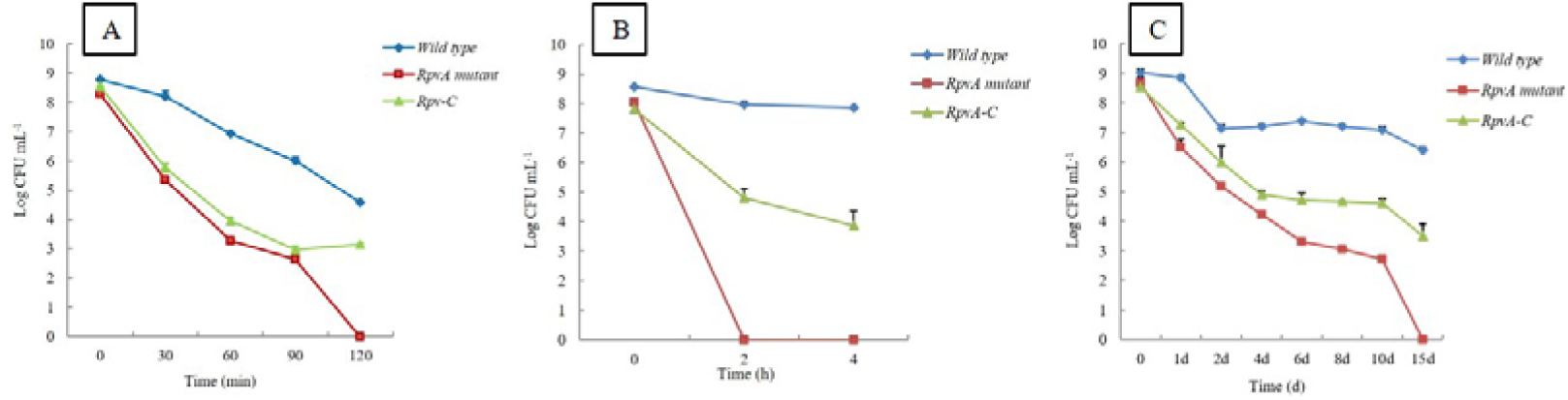
Susceptibilities of the *S. aureus* Newman wild type, *rpvA* mutant, and its complemented strain to stress. A. Susceptibility to heat (58 ± 0.1 °C). B. Susceptibility to H_2_O_2_ (100 mM). C. Susceptibility to starvation in saline.

### rpvA mutant was less able to survive in THP-1 macrophages

The intracellular survival of the *S. aureus* Newman wild type strain, the *rpvA* mutant and its complemented strain was assessed by using human THP-1 phagocytosis test. The survival of the *rpvA* mutant in THP-1 cells decreased 100-fold after it was phagocytized 24 hours, whereas the wild type only decreased 10-fold. The *rpvA* mutant complemented strain restored its survival partially (Figure 4).

**Figure 4.**
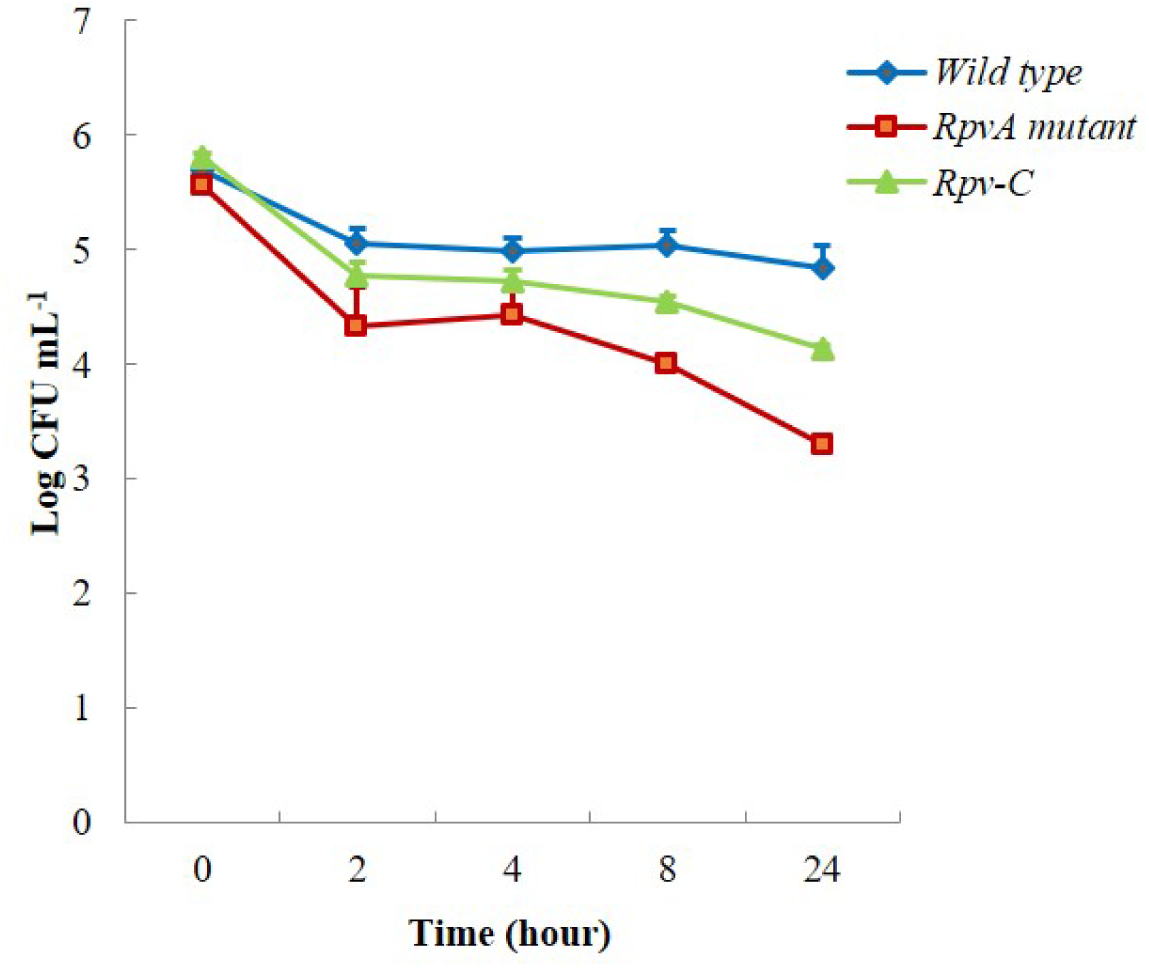
The intracellular anti-phagocytosis of the *S. aureus* Newman wild type, *rpvA* mutant and its complemented strain assessed by human THP-1 monocytes phagocytosis test.

### Virulence of rpvA mutant is severely attenuated in mice

The LD_50_ of the *rpvA* mutant in BALB/c mice was significantly higher than that of the parent strain. No mice died when 8.33 × 10^10^ CFU/mL of the *rpvA* mutant were injected intraperitoneally, indicating that the LD_50_ (log_10_ CFU) of the *rpvA* mutant in BALB/c mice was over 10.921, whereas the LD_50_ of the wild type and the complemented strain were 9.382 and 10.063, respectively.

### Mouse subcutaneous abscess model of rpvA mutant

Mouse model was used to detect the possible difference between *S. aureus* Newman wild type and the *rpvA* mutant in terms of their ability to form skin abscesses. The bacteria reliably formed abscesses and the mice local skin injected with wild type, *rpvA* mutant and its complemented strain all showed swelling after 48 hours following injection. However, the lesion of the mouse skin injected with the *rpvA* mutant was relatively minor, whereas the skin lesion injected with wild strain and the complemented strain displayed a more clear skin ulceration and damage (Figure 5 A). Examination of the HE stained mouse skin lesions discovered that except for the control group, the wild type group, *rpvA* mutant group, and its complemented strain group mice skin lesions all presented as pyogenic inflammation characteristic (Figure 5 B). The wild type group lesions presented both as acute inflammation and skin ulceration (Figure 5 B2, arrow). The *rpvA* mutant group skin lesions presented as acute inflammation mainly but rare skin ulceration (Figure 5 B4). Complemented with *rpvA* restored the mutant ability to cause lesion in the mouse skin (Figure 5 B3, arrow). The bacteria in abscesses were harvested and the numbers were counted as described (47). The Log CFU mL^−1^ of the *rpvA* mutant in the lesions was 3.09 ± 0.12, which was decreased about 10,000-fold compared with the initial injected number. The Log CFU mL^−1^ of the wild type in the lesions was 4.51 ± 0.45, which was decreased about 1000-fold than the initial injected number. Complementation with *rpvA* restored the numbers of bacteria in the abscess (Figure 5 C). These findings indicate that *rpvA* plays prominent roles during *S. aureus* Newman skin infection.

**Figure 5.**
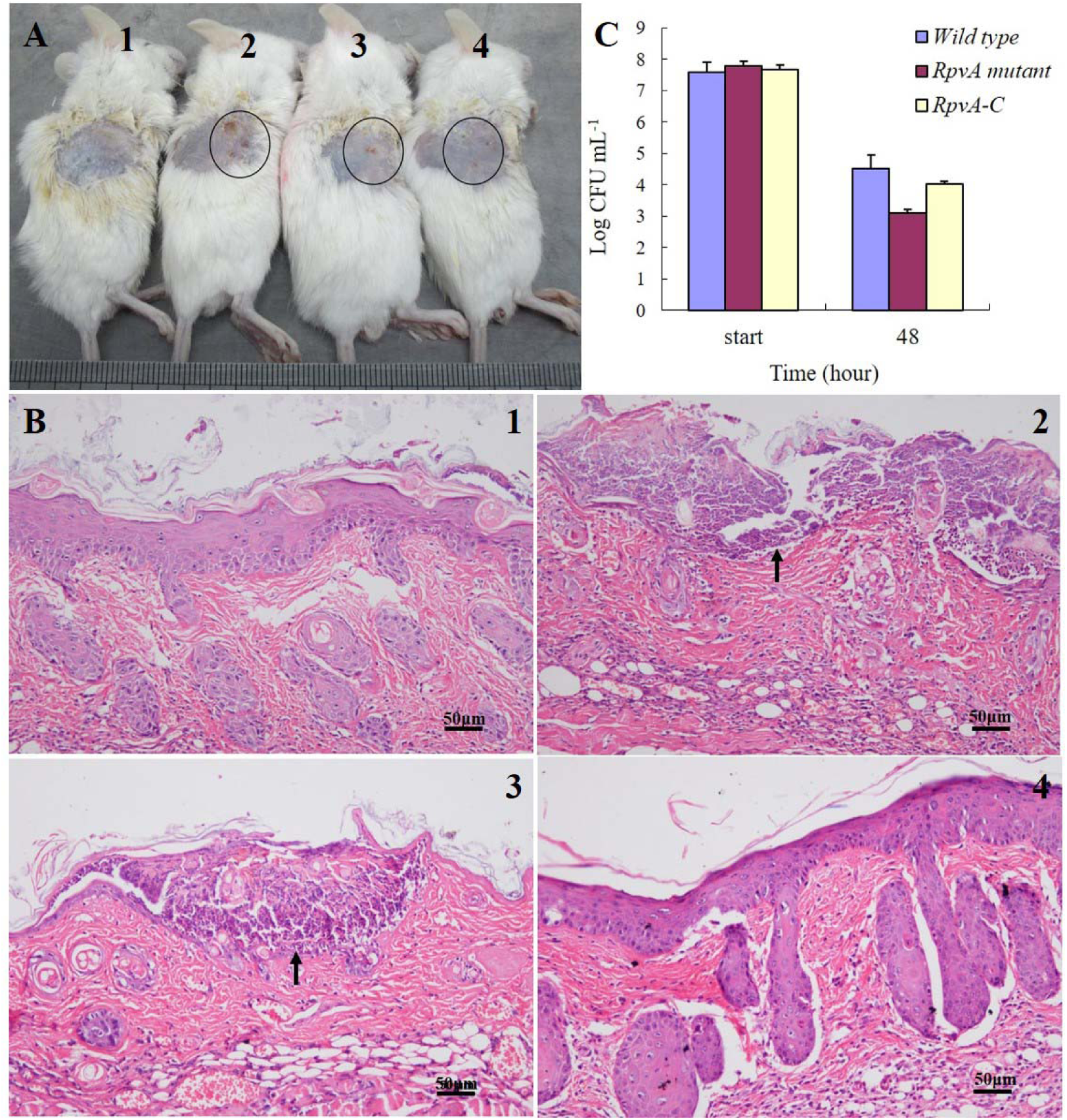
The local skin lesion appearance. A. The injected mice; A1. Control group. The mouse skin was integral and no damage and ulceration. A2. The *S. aureus* Newman strain group. Three obvious damage and ulcers could be seen at the local injection site (circle). A3. The *rpvA* mutant complemented strain group. One obvious damage and ulcer can be seen at the injection site (circle). A4. The *rpvA* mutant group. Only three very slight damage could be seen at the injection site (circle). B. Pathological changes of mice skin lesions; B1. Intact skin of control group mice. B2. Wild type group mice skin. There was obviously acute inflammation of subcutaneous and skin ulceration (arrow). B3. *rpvA* mutant complemented group. There was obviously acute inflammation of subcutaneous and skin ulceration (arrow). B4. *rpvA* mutant group. There was obviously acute inflammation of subcutaneous, whereas no skin ulceration could be seen. C. Number of viable bacteria counted in mice subcutaneous abscess.

### RNAseq demonstrates global regulatory impact of RpvA

RNAseq results revealed that in 6-hour cultures, 38 genes in the *rpvA* mutant were down-regulated, while 73 genes were up-regulated significantly (log2.fold_change.>2) compared with the parent strain (Table 4). qPCR confirmed that except *pgk*, all other genes (*eno*, *pgm*, *tpi*, *gapA*, *mtlD*, *crtM*, *hla*, and *hlgB*) expression trends were similar to those from the RNA-seq results. These significant differentially expressed genes are distributed in metabolism pathways, including global and overview metabolism, carbohydrate metabolism, energy metabolism, lipid metabolism, nucleotide metabolism, amino acid metabolism, metabolism of cofactors and vitamins and others, genetic information processing, environmental information processing, and virulence (Figure 6 and 7). Interestingly, the expression levels of genes in the *rpvA* mutant involved in carbohydrate metabolism (*eno*, *pgm*, *tpi*, *gapA*, *glpD*, *acs*, *etc*.) and amino acid biosynthesis and metabolism (*glmS*, *pgm*, *narJ*, *narG*, *narH*, *alsS*, *eno*, *acs*, etc.) were significantly up-regulated, while genes involved in virulence (*hla*, *hlgB*, *crtI*, *crtN*, *crtM*, *set1nm*) were significantly down-regulated compared to the wild type strain. These results suggest that the *rpvA* mutant is more active in metabolism than the wild type strain, while the wild strain is more virulent under the same conditions than the *rpvA* mutant.

**Figure 6.**
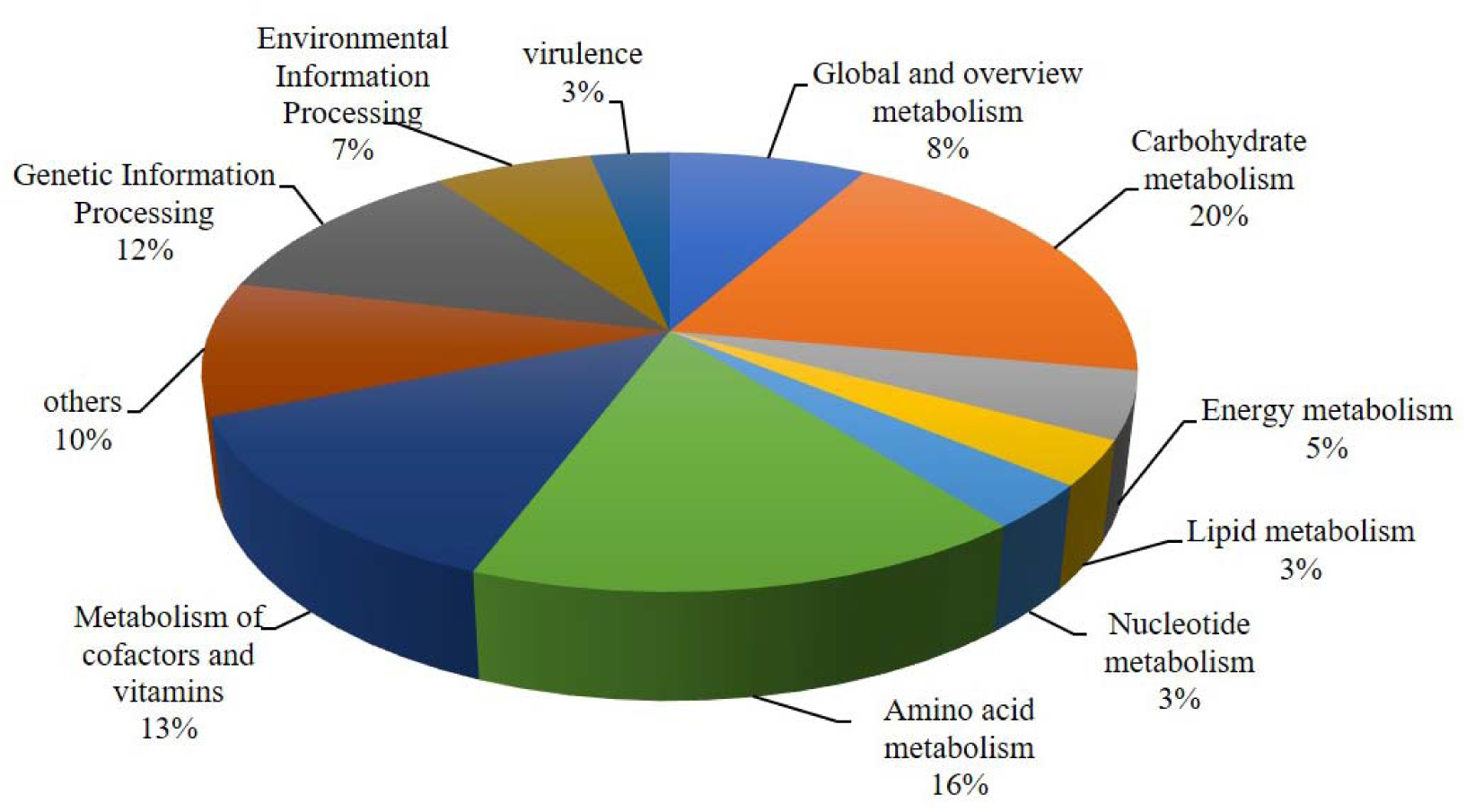
Pathway ration distribution map of differentially expressed genes regulated by RpvA.

**Figure 7.**
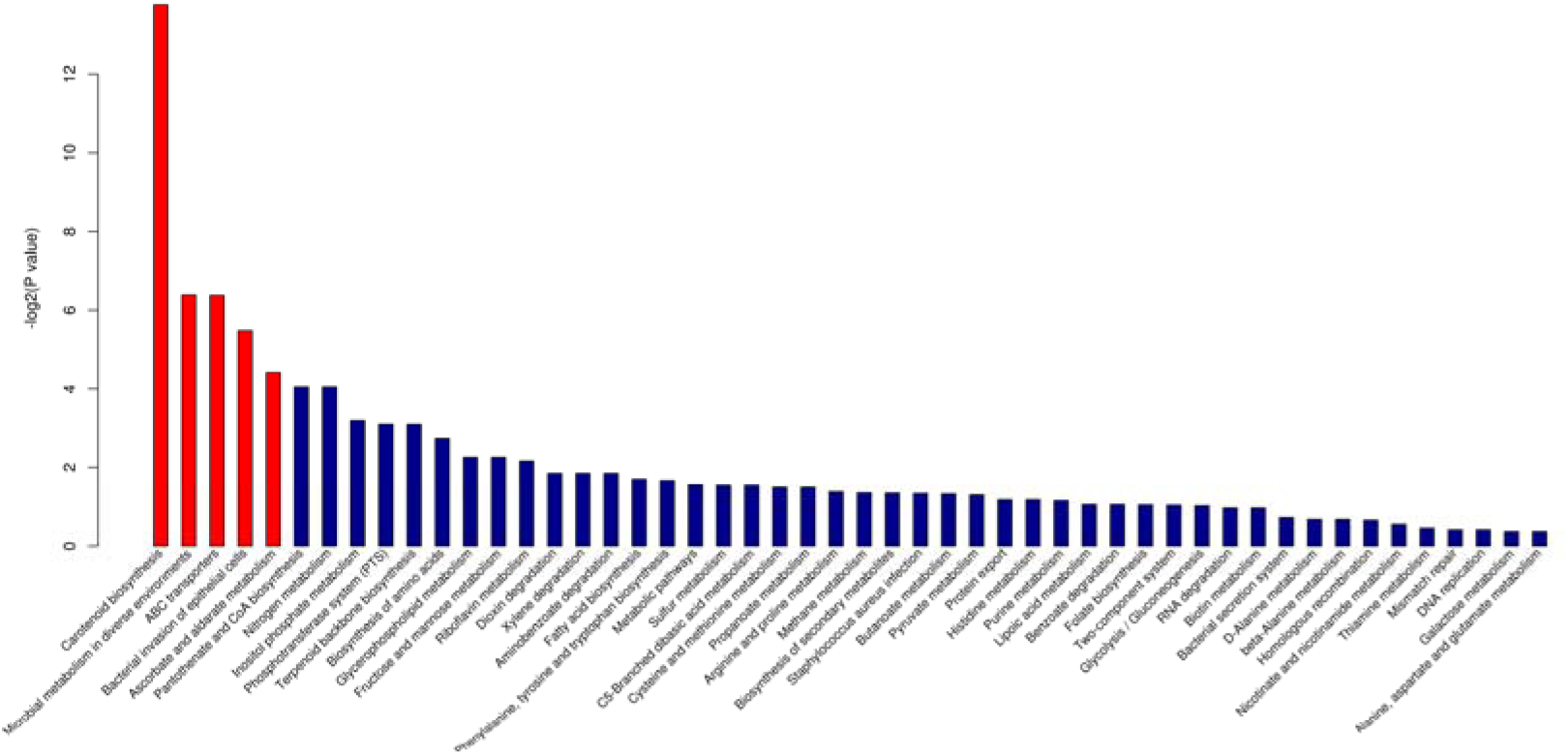
The KEGG pathway annotation *P*-value distribution of transcriptome (*P*-value < 0.05 in red bar pathway)

**Table 4.**
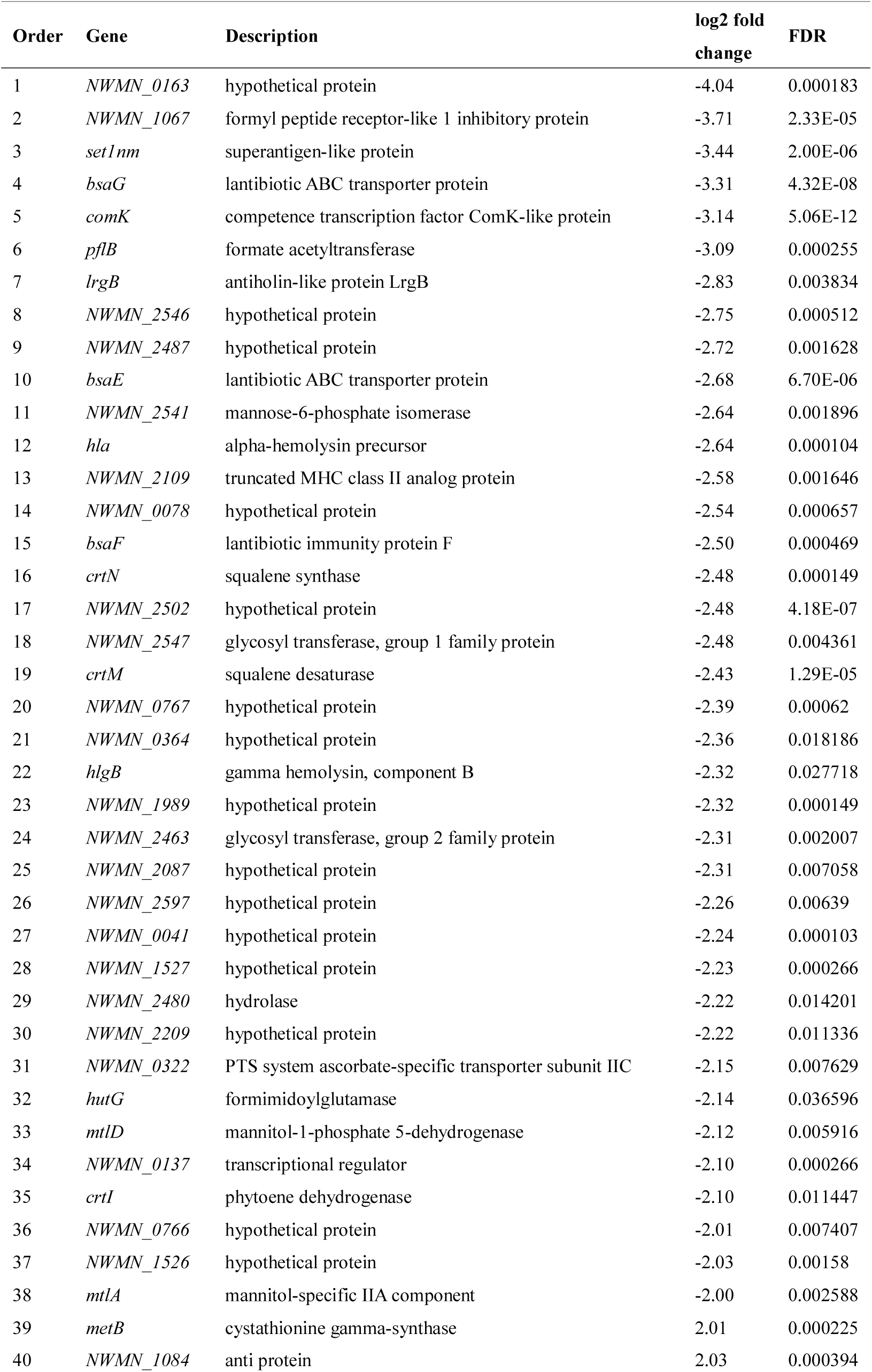
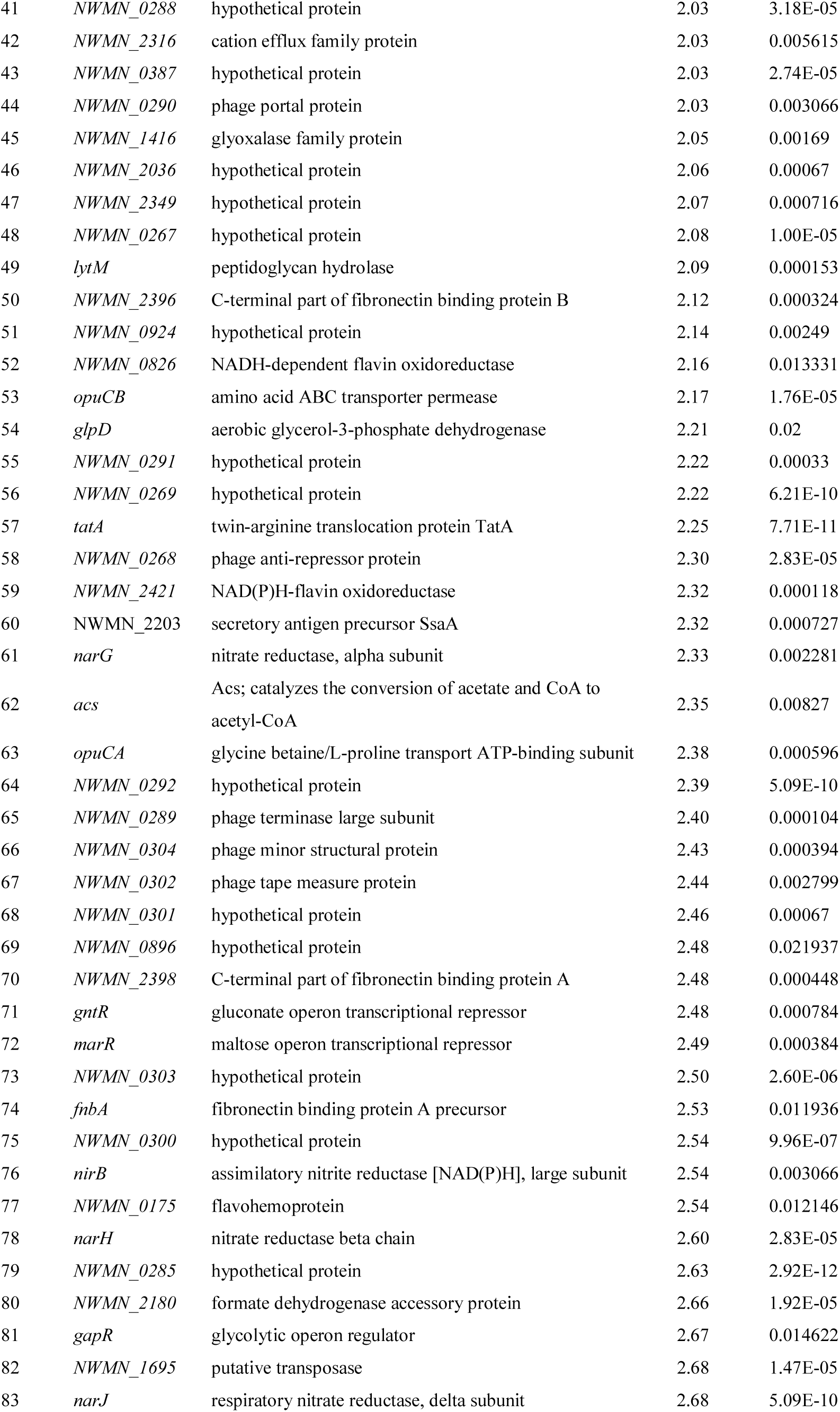
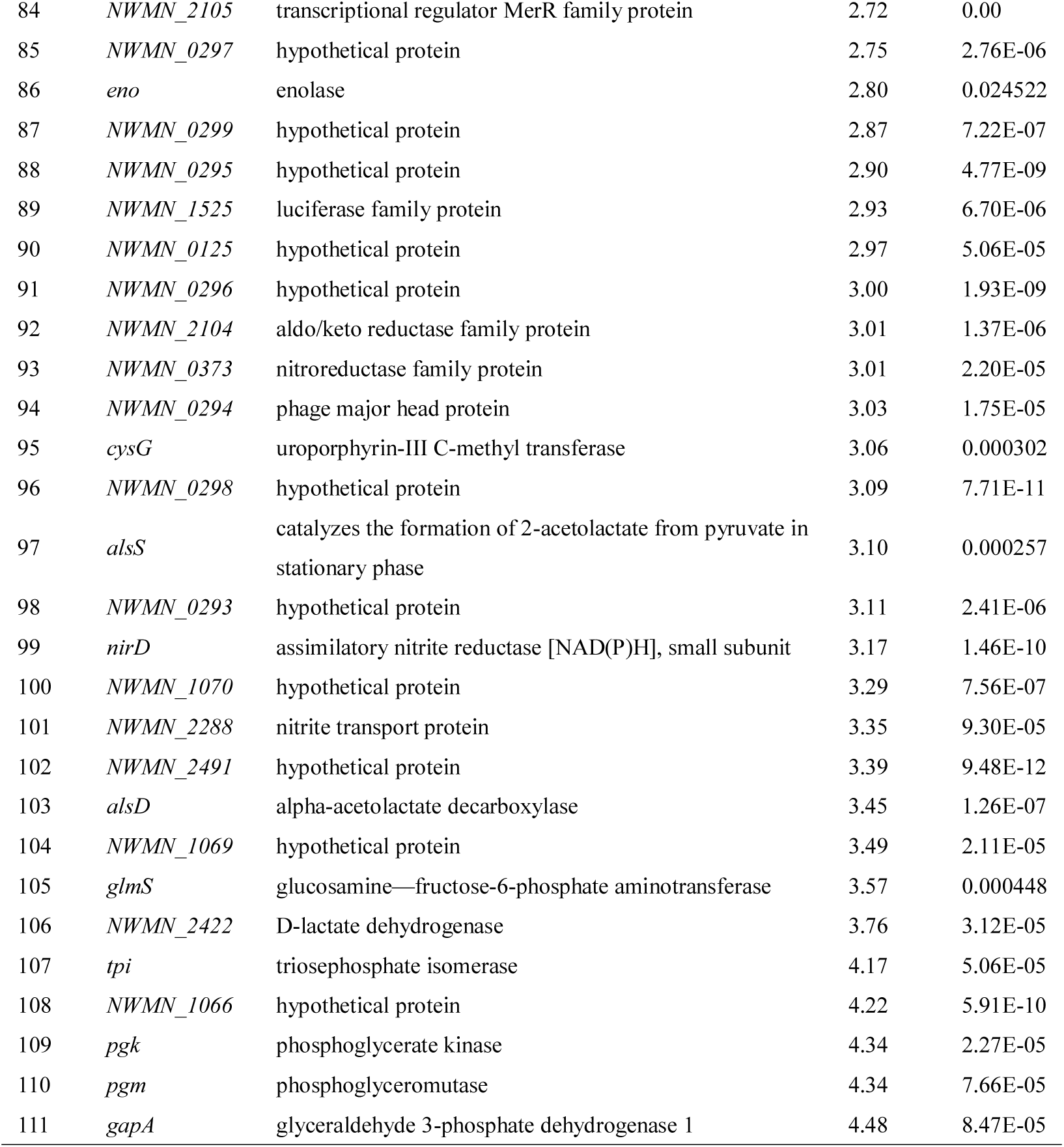
The differentially expressed genes between *rpvA* mutant and wild type after 6 hours incubation

| Order | Gene | Description | log2 fold change | FDR |
| --- | --- | --- | --- | --- |
| 1 | *NWMN_0163* | hypothetical protein | -4.04 | 0.000183 |
| 2 | *NWMN_1067* | formyl peptide receptor-like 1 inhibitory protein | -3.71 | 2.33E-05 |
| 3 | *set1nm* | superantigen-like protein | -3.44 | 2.00E-06 |
| 4 | *bsaG* | lantibiotic ABC transporter protein | -3.31 | 4.32E-08 |
| 5 | *comK* | competence transcription factor ComK-like protein | -3.14 | 5.06E-12 |
| 6 | *pflB* | formate acetyltransferase | -3.09 | 0.000255 |
| 7 | *lrgB* | antiholin-like protein LrgB | -2.83 | 0.003834 |
| 8 | *NWMN_2546* | hypothetical protein | -2.75 | 0.000512 |
| 9 | *NWMN_2487* | hypothetical protein | -2.72 | 0.001628 |
| 10 | *bsaE* | lantibiotic ABC transporter protein | -2.68 | 6.70E-06 |
| 11 | *NWMN_2541* | mannose-6-phosphate isomerase | -2.64 | 0.001896 |
| 12 | *hla* | alpha-hemolysin precursor | -2.64 | 0.000104 |
| 13 | *NWMN_2109* | truncated MHC class II analog protein | -2.58 | 0.001646 |
| 14 | *NWMN_0078* | hypothetical protein | -2.54 | 0.000657 |
| 15 | *bsaF* | lantibiotic immunity protein F | -2.50 | 0.000469 |
| 16 | *crtN* | squalene synthase | -2.48 | 0.000149 |
| 17 | *NWMN_2502* | hypothetical protein | -2.48 | 4.18E-07 |
| 18 | *NWMN_2547* | glycosyl transferase, group 1 family protein | -2.48 | 0.004361 |
| 19 | *crtM* | squalene desaturase | -2.43 | 1.29E-05 |
| 20 | *NWMN_0767* | hypothetical protein | -2.39 | 0.00062 |
| 21 | *NWMN_0364* | hypothetical protein | -2.36 | 0.018186 |
| 22 | *hlgB* | gamma hemolysin, component B | -2.32 | 0.027718 |
| 23 | *NWMN_1989* | hypothetical protein | -2.32 | 0.000149 |
| 24 | *NWMN_2463* | glycosyl transferase, group 2 family protein | -2.31 | 0.002007 |
| 25 | *NWMN_2087* | hypothetical protein | -2.31 | 0.007058 |
| 26 | *NWMN_2597* | hypothetical protein | -2.26 | 0.00639 |
| 27 | *NWMN_0041* | hypothetical protein | -2.24 | 0.000103 |
| 28 | *NWMN_1527* | hypothetical protein | -2.23 | 0.000266 |
| 29 | *NWMN_2480* | hydrolase | -2.22 | 0.014201 |
| 30 | *NWMN_2209* | hypothetical protein | -2.22 | 0.011336 |
| 31 | *NWMN_0322* | PTS system ascorbate-specific transporter subunit IIC | -2.15 | 0.007629 |
| 32 | *hutG* | formimidoylglutamase | -2.14 | 0.036596 |
| 33 | *mtlD* | mannitol-1-phosphate 5-dehydrogenase | -2.12 | 0.005916 |
| 34 | *NWMN_0137* | transcriptional regulator | -2.10 | 0.000266 |
| 35 | *crtI* | phytoene dehydrogenase | -2.10 | 0.011447 |
| 36 | *NWMN_0766* | hypothetical protein | -2.01 | 0.007407 |
| 37 | *NWMN_1526* | hypothetical protein | -2.03 | 0.00158 |
| 38 | *mtlA* | mannitol-specific IIA component | -2.00 | 0.002588 |
| 39 | *metB* | cystathionine gamma-synthase | 2.01 | 0.000225 |
| 40 | *NWMN_1084* | anti protein | 2.03 | 0.000394 |
| 41 | *NWMN_0288* | hypothetical protein | 2.03 | 3.18E-05 |
| 42 | *NWMN_2316* | cation efflux family protein | 2.03 | 0.005615 |
| 43 | *NWMN_0387* | hypothetical protein | 2.03 | 2.74E-05 |
| 44 | *NWMN_0290* | phage portal protein | 2.03 | 0.003066 |
| 45 | *NWMN_1416* | glyoxalase family protein | 2.05 | 0.00169 |
| 46 | *NWMN_2036* | hypothetical protein | 2.06 | 0.00067 |
| 47 | *NWMN_2349* | hypothetical protein | 2.07 | 0.000716 |
| 48 | *NWMN_0267* | hypothetical protein | 2.08 | 1.00E-05 |
| 49 | *lytM* | peptidoglycan hydrolase | 2.09 | 0.000153 |
| 50 | *NWMN_2396* | C-terminal part of fibronectin binding protein B | 2.12 | 0.000324 |
| 51 | *NWMN_0924* | hypothetical protein | 2.14 | 0.00249 |
| 52 | *NWMN_0826* | NADH-dependent flavin oxidoreductase | 2.16 | 0.013331 |
| 53 | *opuCB* | amino acid ABC transporter permease | 2.17 | 1.76E-05 |
| 54 | *glpD* | aerobic glycerol-3-phosphate dehydrogenase | 2.21 | 0.02 |
| 55 | *NWMN_0291* | hypothetical protein | 2.22 | 0.00033 |
| 56 | *NWMN_0269* | hypothetical protein | 2.22 | 6.21E-10 |
| 57 | *tatA* | twin-arginine translocation protein TatA | 2.25 | 7.71E-11 |
| 58 | *NWMN_0268* | phage anti-repressor protein | 2.30 | 2.83E-05 |
| 59 | *NWMN_2421* | NAD(P)H-flavin oxidoreductase | 2.32 | 0.000118 |
| 60 | *NWMN_2203* | secretory antigen precursor SsaA | 2.32 | 0.000727 |
| 61 | *narG* | nitrate reductase, alpha subunit | 2.33 | 0.002281 |
| 62 | *acs* | Acs; catalyzes the conversion of acetate and CoA to acetyl-CoA | 2.35 | 0.00827 |
| 63 | *opuCA* | glycine betaine/L-proline transport ATP-binding subunit | 2.38 | 0.000596 |
| 64 | *NWMN_0292* | hypothetical protein | 2.39 | 5.09E-10 |
| 65 | *NWMN_0289* | phage terminase large subunit | 2.40 | 0.000104 |
| 66 | *NWMN_0304* | phage minor structural protein | 2.43 | 0.000394 |
| 67 | *NWMN_0302* | phage tape measure protein | 2.44 | 0.002799 |
| 68 | *NWMN_0301* | hypothetical protein | 2.46 | 0.00067 |
| 69 | *NWMN_0896* | hypothetical protein | 2.48 | 0.021937 |
| 70 | *NWMN_2398* | C-terminal part of fibronectin binding protein A | 2.48 | 0.000448 |
| 71 | *gntR* | gluconate operon transcriptional repressor | 2.48 | 0.000784 |
| 72 | *marR* | maltose operon transcriptional repressor | 2.49 | 0.000384 |
| 73 | *NWMN_0303* | hypothetical protein | 2.50 | 2.60E-06 |
| 74 | *fnbA* | fibronectin binding protein A precursor | 2.53 | 0.011936 |
| 75 | *NWMN_0300* | hypothetical protein | 2.54 | 9.96E-07 |
| 76 | *nirB* | assimilatory nitrite reductase [NAD(P)H], large subunit | 2.54 | 0.003066 |
| 77 | *NWMN_0175* | flavohemoprotein | 2.54 | 0.012146 |
| 78 | *narH* | nitrate reductase beta chain | 2.60 | 2.83E-05 |
| 79 | *NWMN_0285* | hypothetical protein | 2.63 | 2.92E-12 |
| 80 | *NWMN_2180* | formate dehydrogenase accessory protein | 2.66 | 1.92E-05 |
| 81 | *gapR* | glycolytic operon regulator | 2.67 | 0.014622 |
| 82 | *NWMN_1695* | putative transposase | 2.68 | 1.47E-05 |
| 83 | *narJ* | respiratory nitrate reductase, delta subunit | 2.68 | 5.09E-10 |
| 84 | *NWMN_2105* | transcriptional regulator MerR family protein | 2.72 | 0.00 |
| 85 | *NWMN_0297* | hypothetical protein | 2.75 | 2.76E-06 |
| 86 | *eno* | enolase | 2.80 | 0.024522 |
| 87 | *NWMN_0299* | hypothetical protein | 2.87 | 7.22E-07 |
| 88 | *NWMN_0295* | hypothetical protein | 2.90 | 4.77E-09 |
| 89 | *NWMN_1525* | luciferase family protein | 2.93 | 6.70E-06 |
| 90 | *NWMN_0125* | hypothetical protein | 2.97 | 5.06E-05 |
| 91 | *NWMN_0296* | hypothetical protein | 3.00 | 1.93E-09 |
| 92 | *NWMN_2104* | aldo/keto reductase family protein | 3.01 | 1.37E-06 |
| 93 | *NWMN_0373* | nitroreductase family protein | 3.01 | 2.20E-05 |
| 94 | *NWMN_0294* | phage major head protein | 3.03 | 1.75E-05 |
| 95 | *cysG* | uroporphyrin-III C-methyl transferase | 3.06 | 0.000302 |
| 96 | *NWMN_0298* | hypothetical protein | 3.09 | 7.71E-11 |
| 97 | *alsS* | catalyzes the formation of 2-acetolactate from pyruvate in stationary phase | 3.10 | 0.000257 |
| 98 | *NWMN_0293* | hypothetical protein | 3.11 | 2.41E-06 |
| 99 | *nirD* | assimilatory nitrite reductase [NAD(P)H], small subunit | 3.17 | 1.46E-10 |
| 100 | *NWMN_1070* | hypothetical protein | 3.29 | 7.56E-07 |
| 101 | *NWMN_2288* | nitrite transport protein | 3.35 | 9.30E-05 |
| 102 | *NWMN_2491* | hypothetical protein | 3.39 | 9.48E-12 |
| 103 | *alsD* | alpha-acetolactate decarboxylase | 3.45 | 1.26E-07 |
| 104 | *NWMN_1069* | hypothetical protein | 3.49 | 2.11E-05 |
| 105 | *glmS* | glucosamine—fructose-6-phosphate aminotransferase | 3.57 | 0.000448 |
| 106 | *NWMN_2422* | D-lactate dehydrogenase | 3.76 | 3.12E-05 |
| 107 | *tpi* | triosephosphate isomerase | 4.17 | 5.06E-05 |
| 108 | *NWMN_1066* | hypothetical protein | 4.22 | 5.91E-10 |
| 109 | *pgk* | phosphoglycerate kinase | 4.34 | 2.27E-05 |
| 110 | *pgm* | phosphoglyceromutase | 4.34 | 7.66E-05 |
| 111 | *gapA* | glyceraldehyde 3-phosphate dehydrogenase 1 | 4.48 | 8.47E-05 |

### Metabolomics analysis show changes in metabolites after rpvA mutation

Both unsupervised and supervised multivariate analyses including PCA, PLS, OPLS-DA were performed to further evaluate metabolic perturbations resulting from the *rpvA* mutation in 6-hour cultures. PCA provided a global overview of variance between wild type (WT) and *rpvA* mutant groups. The PCA score plot in both ESI positive and negative modes distinguished clearly between samples from these two groups (Figure 8 A and B). In the supervised OPLS-DA models, it became evident that the wild type group was clearly distinct from the *rpvA* mutant group, and the R2 and Q2 values of the OPLS-DA models parameters showed good adaptability and high predictability in ESI^+^ and ESI^−^ modes (Figure 8 C, D). Permutation tests showed a high significance and confidence for further identification of the distinct metabolites (Figure 8 E, F). A total of 76 differential metabolites were identified in 6-hour cultures between the *rpvA* mutant and wild type samples, of which 70 were decreased (Fold change < 1) and only 6 were increased (Fold change > 1) (Table 5). These metabolites are distributed in global and overview metabolism, carbohydrate metabolism, energy metabolism, lipid metabolism, nucleotide metabolism, amino acid metabolism, metabolism of cofactors and vitamins and others, genetic information processing, environmental information processing (Figure 9 and 10). Among the decreased metabolites, acetyl-CoA, dihydroxyacetone phosphate, phosphoenolpyruvate, L-glutamate, L-aspartate, L-glutamine, DL-lactate, L-lysine, 2-oxoadipic acid, glycerol 3-phosphate, L-phenylalanine, L-tryptophan, and L-arginine *etc*., distributed in several metabolic pathways. Interestingly, citrate is increased in multiple metabolic pathways.

**Figure 8.**
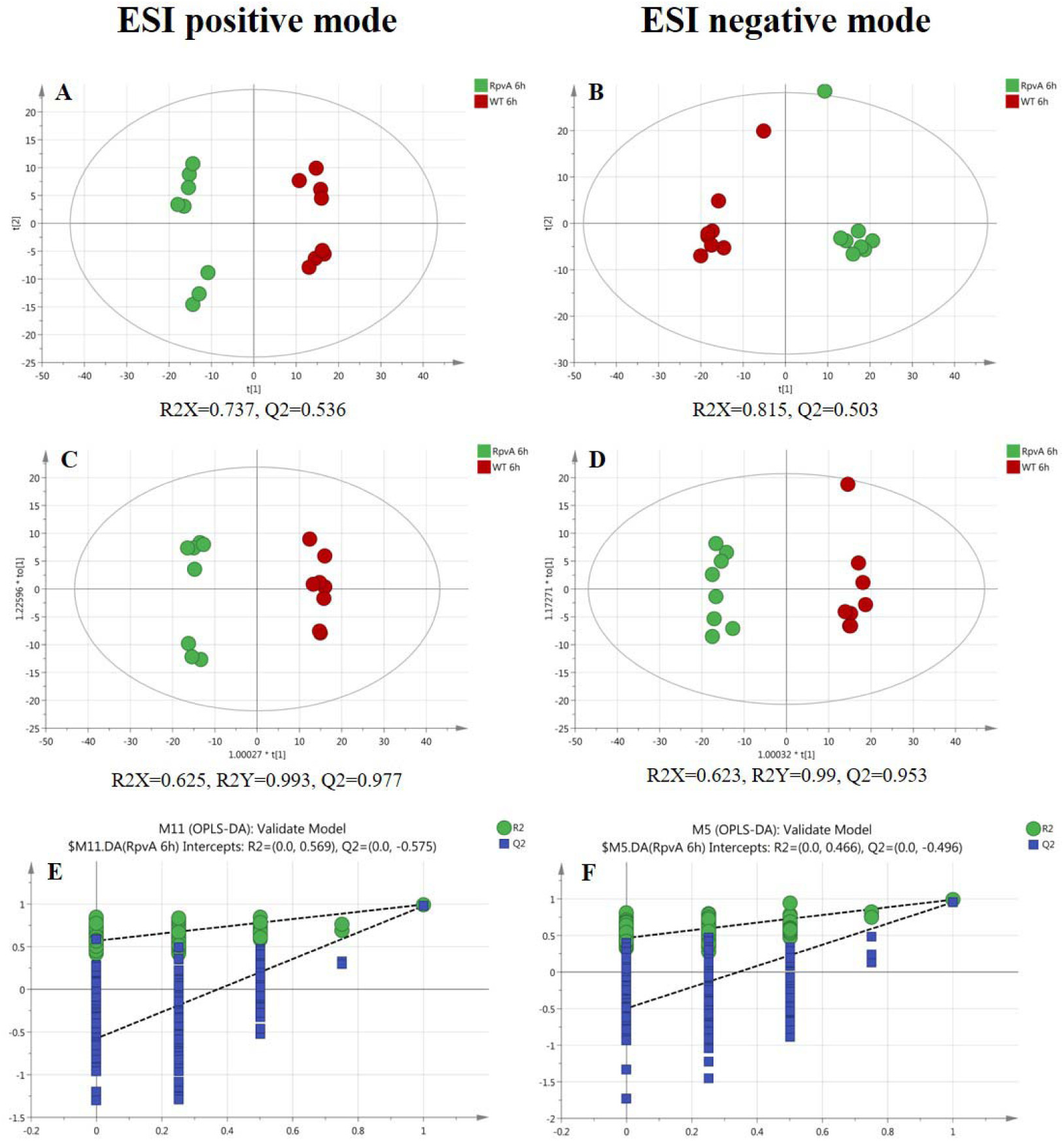
PCA and OPLS-DA score plots in both ESI positive and negative mode. PCA score plot from *rpvA* mutant group vs wild type group in ESI positive mode (A) and negative mode (B). OPLS-DA score plot in each group in ESI positive mode (C) and negative mode (D). The permutation test on the OPLS-DA in ESI positive mode (E) and negative mode (F).

**Figure 9.**
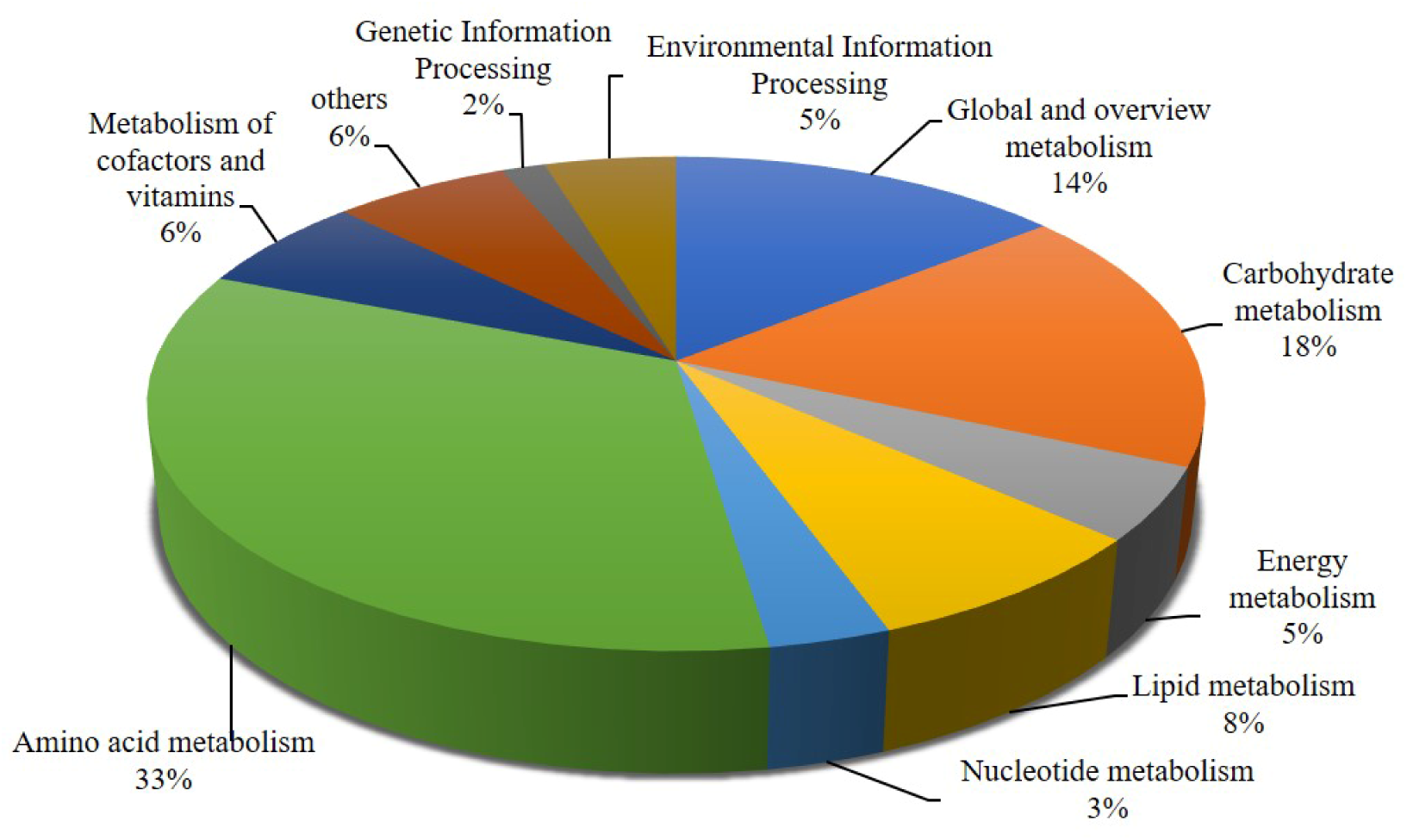
Pathway distribution map of differential metabolites regulated by RpvA

**Figure 10.**
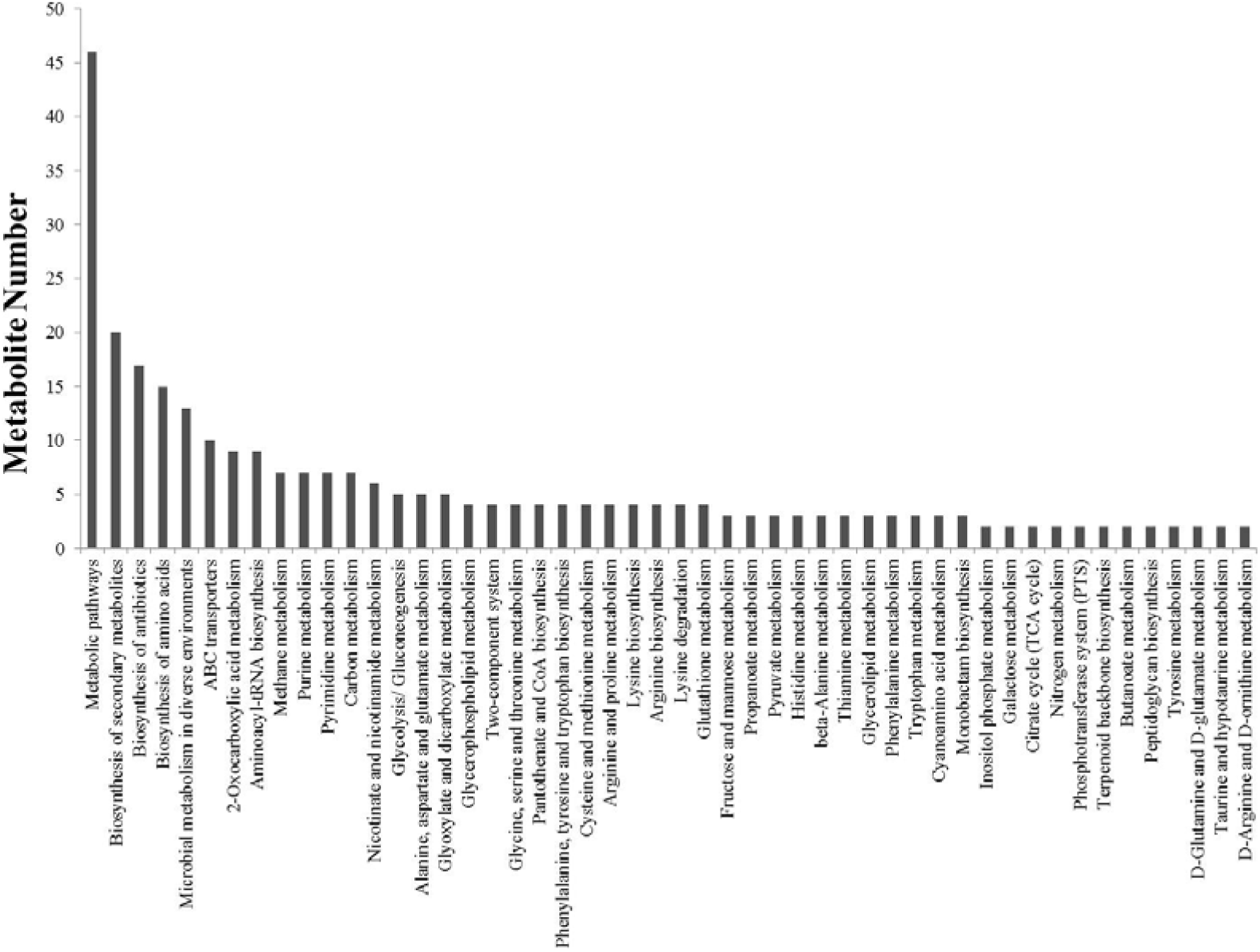
KEGG pathways where differential metabolites regulated by RpvA are involved.

**Table 5.**
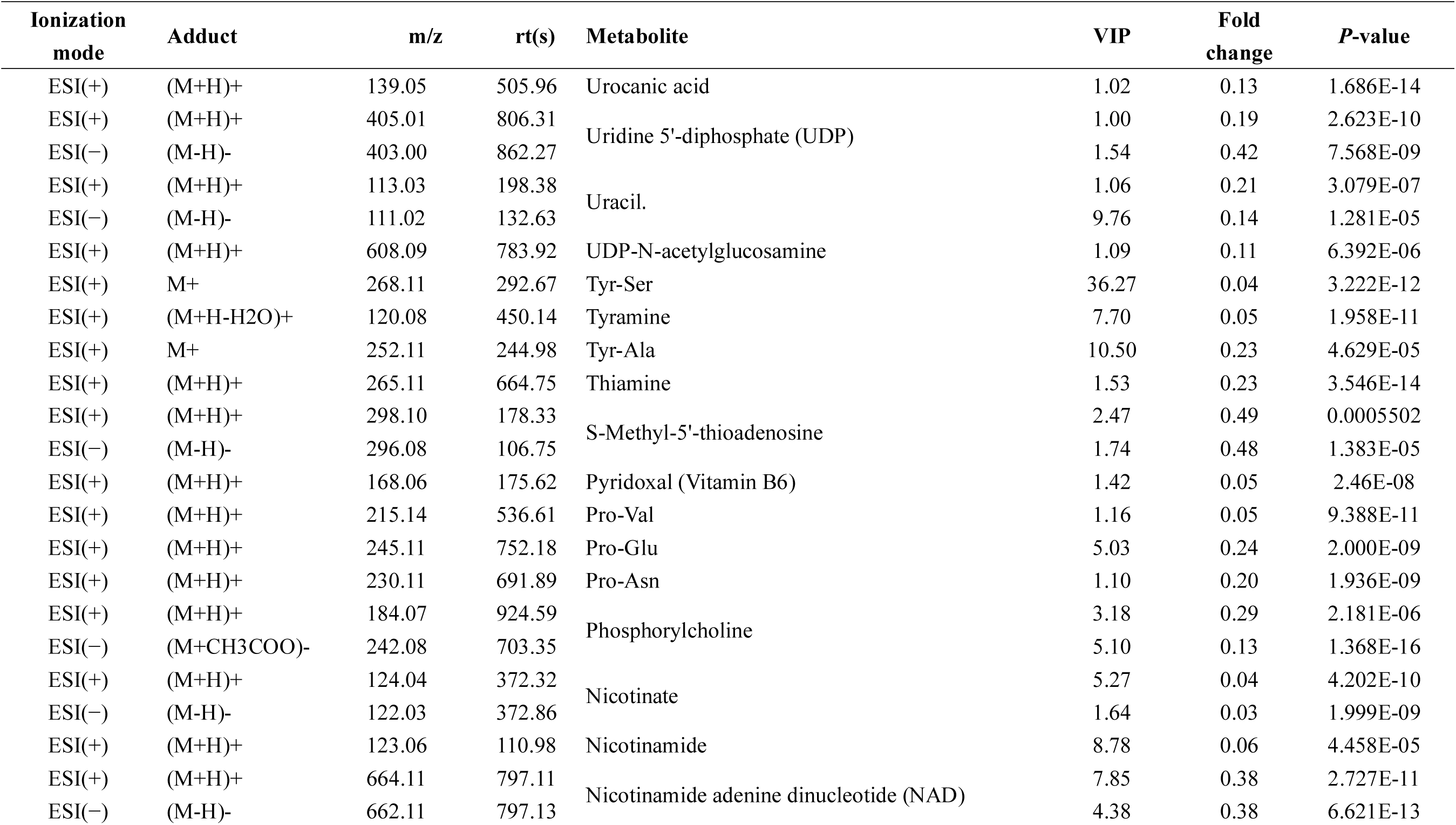
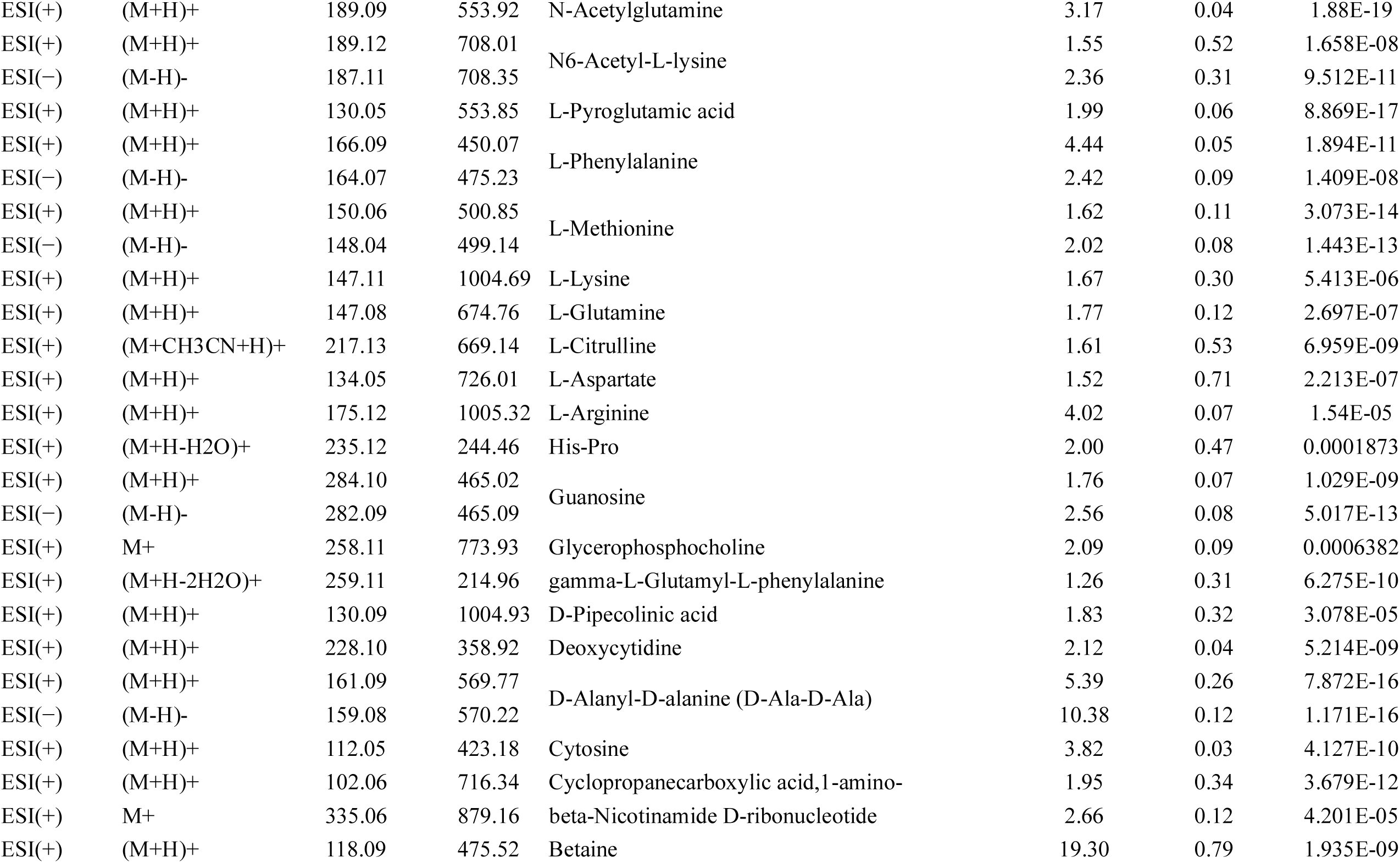
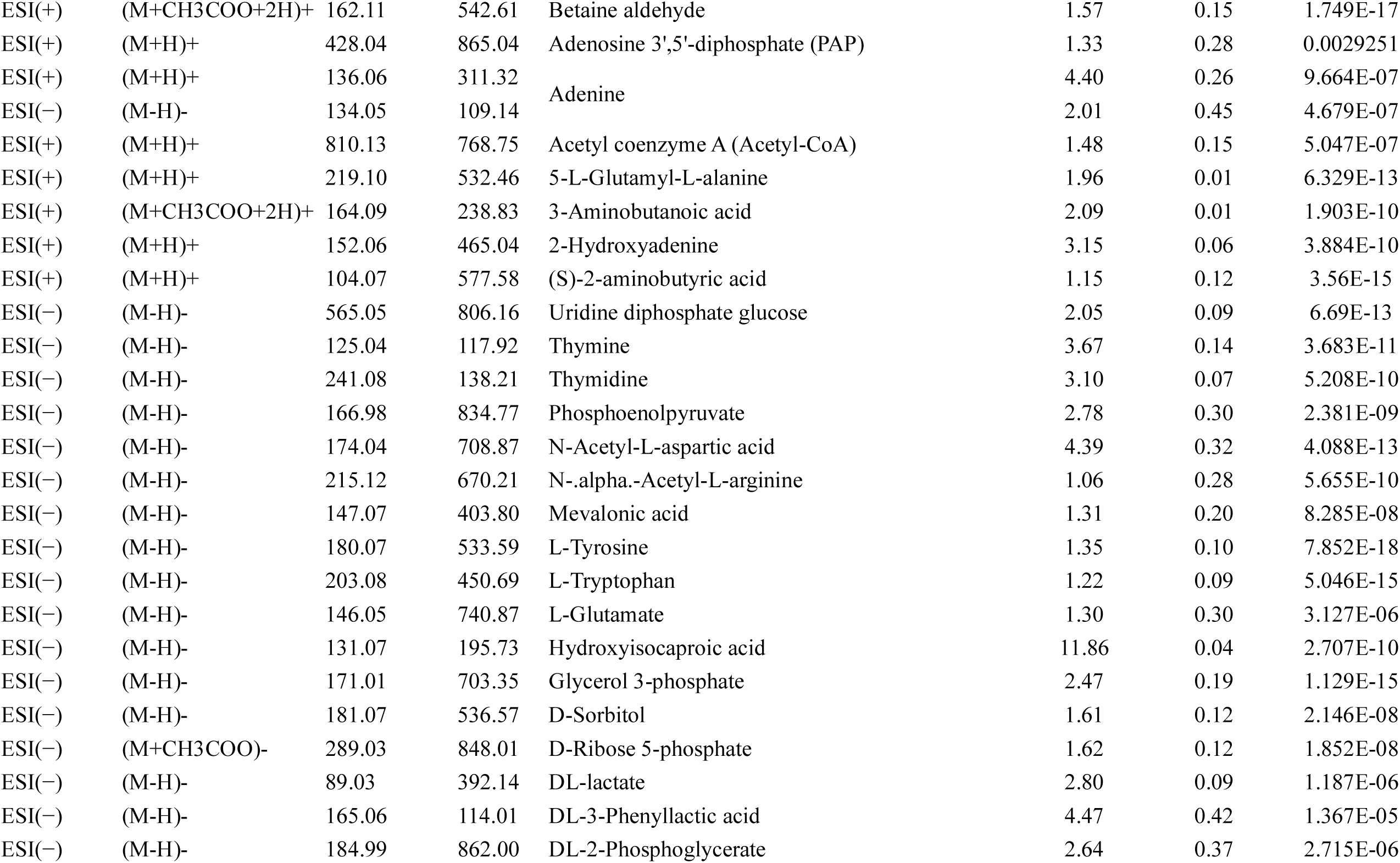
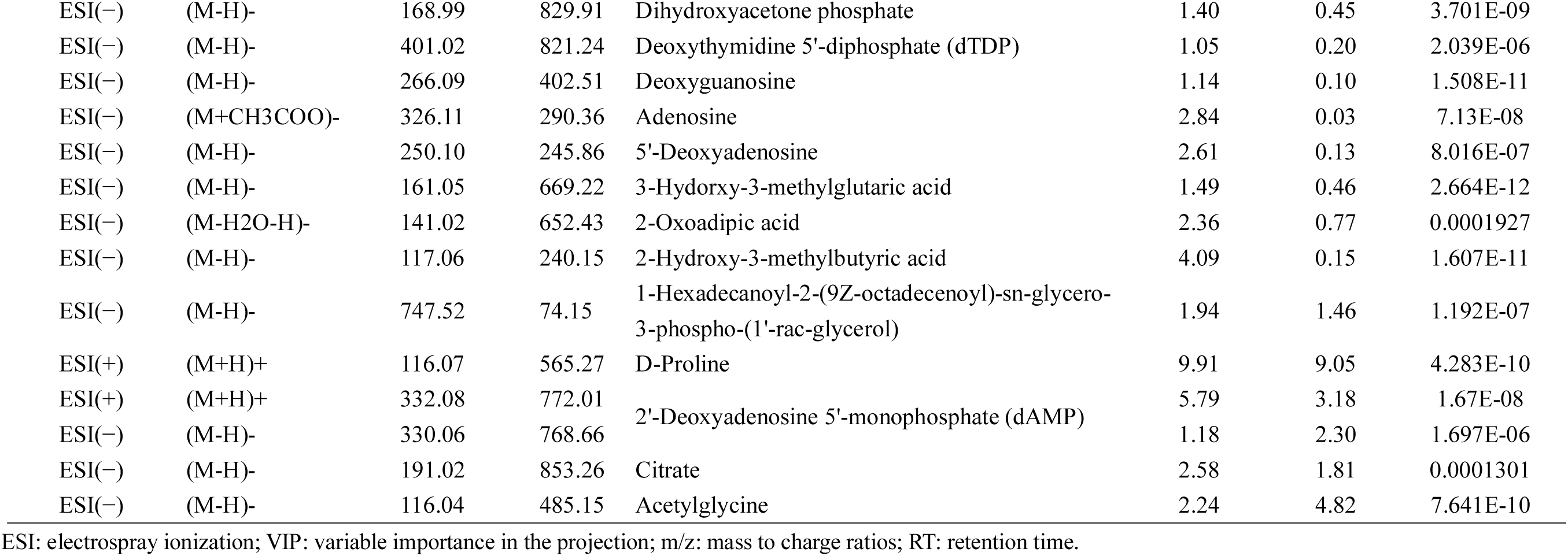
Differential metabolites between *S. aureus rpvA* mutant and wild type after 6 hours incubation

### Transcriptome-metabolome data co-analysis

To associate the results of transcriptomics and metabolomics analyses, we compared the metabolic pathways distributed by differentially expressed genes and metabolites. In total, matches to 19 metabolic pathways showed changes. Further analysis of these differential metabolic pathways revealed that they were mainly involved in global overall metabolism, including carbohydrate metabolism, amino acid metabolism, energy metabolism, metabolism of cofactors and vitamins, lipid metabolism and signal transductions (Figure 11 A and B).

**Figure 11.**
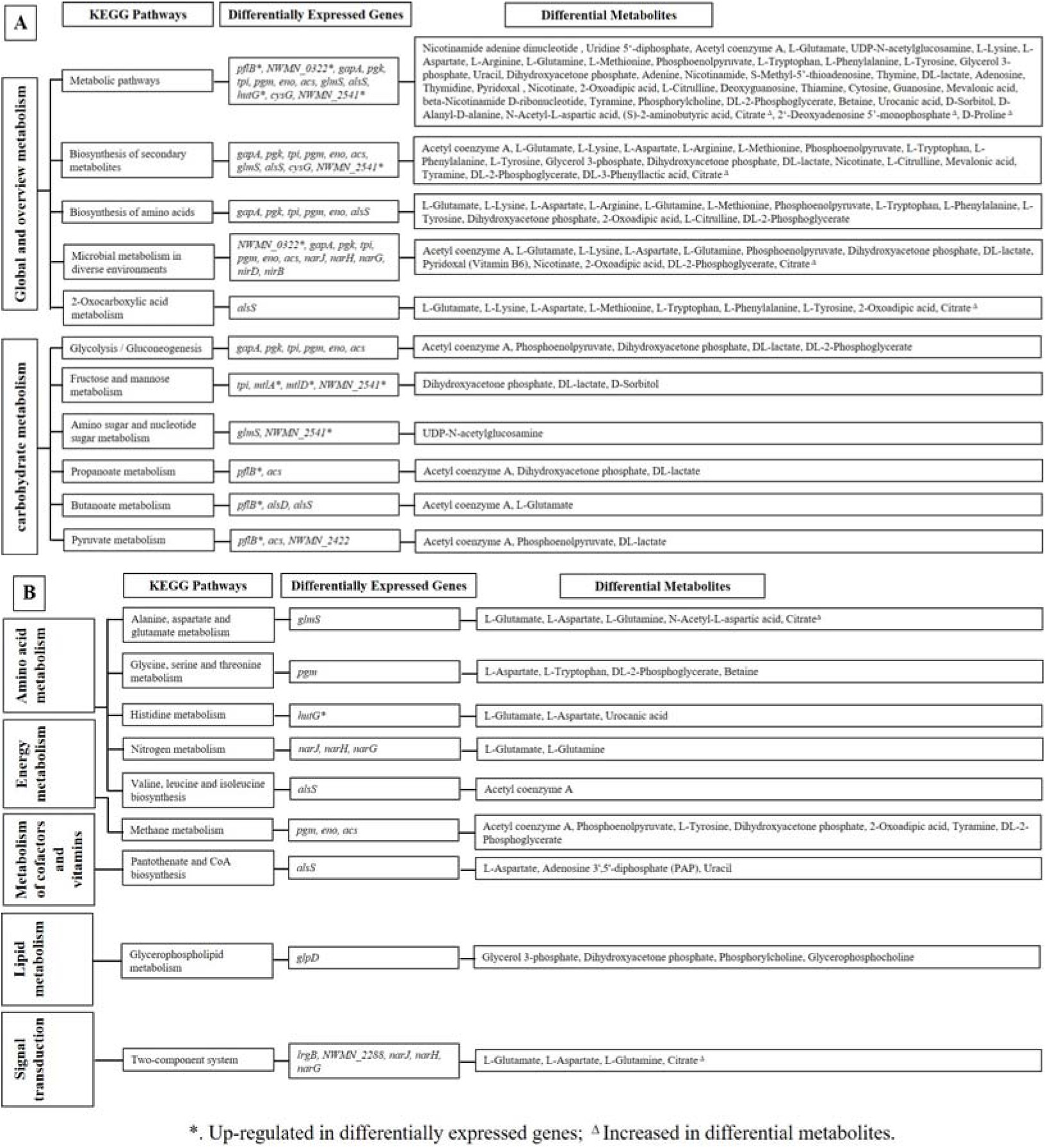
Combined analysis of transcriptome and metabolome of RpvA-mediated metabolism. A. Differential genes and metabolites that are involved in global metabolism and carbohydrate metabolism pathways. B. Differential genes and metabolites that are involved in amino acid metabolism, energy metabolism, metabolism of cofactors and vitamins, lipid metabolism and signal transduction.

### Virulence gene expression of rpvA mutant was significantly down-regulated

In order to analyze the reason for the decreased virulence of the *rpvA* mutation, we performed qPCR to detect the virulence gene expression levels of the *rpvA* mutant and wild type in 6-hour cultures. The results showed that among the main virulence genes of *S. aureus*, except for *lukE*, the expression levels of other virulence genes including *hla*, *hlgA*, *hlgB*, *hlgC*, *lukS*, *lukF*, *lukD*, *sea*, *coa*, and *eta* in the *rpvA* mutant were significantly lower than those in the wild type (*P* value < 0.05) (Figure 12).

**Figure 12.**
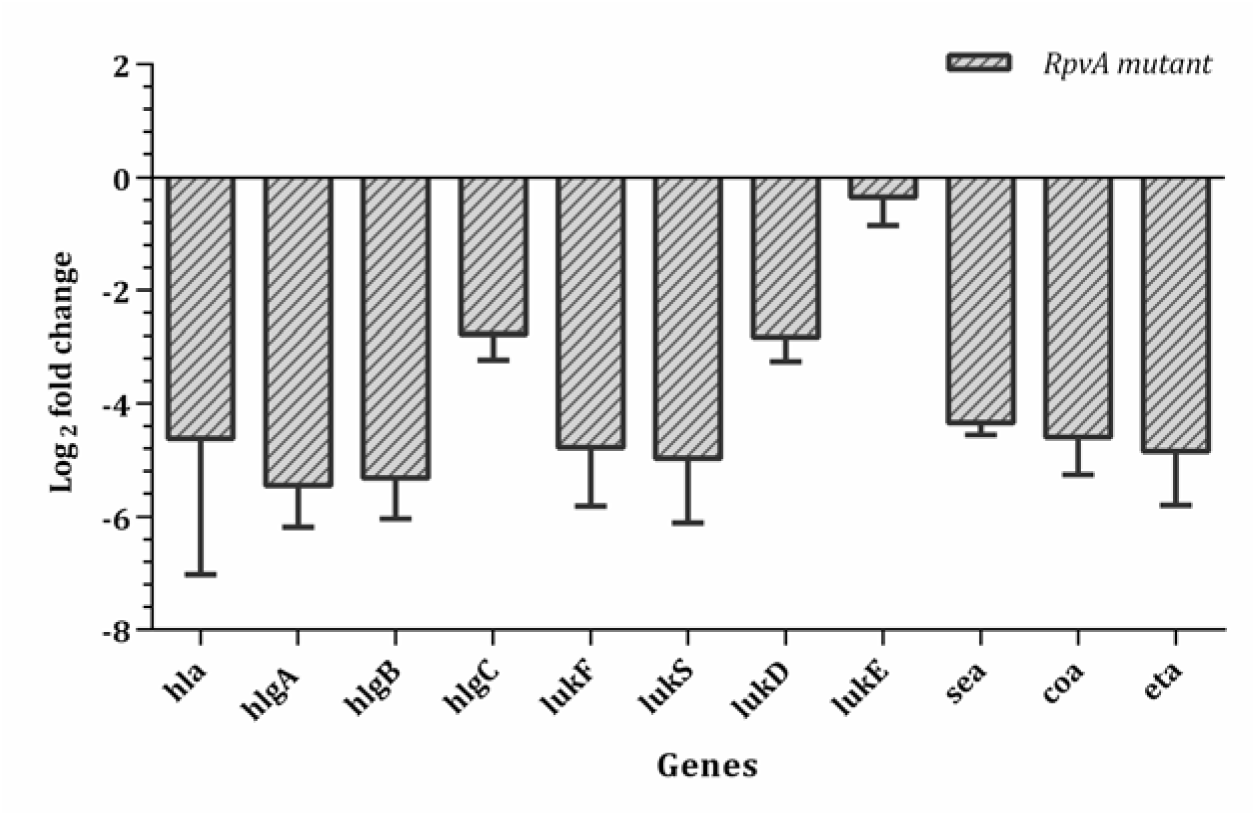
Virulence gene expression levels of *rpvA* mutant against wild type in 6-hour cultures. * *P* value < 0.05.

### *RpvA* fragment can binds to promoters of crtM, hla and hlgB

Due to the complex structure of RpvA, we could only express a soluble truncated portion of GB1-VpmR-S1 in an IPTG inducible *E. coli* C43(DE3) culture (Figure 13 A). Based on our findings on the phenotypes of the *rpvA* mutant and the RNA-seq results, we speculated that the transcriptional regulator RpvA may modulate virulence genes (*hla, hlgB*) and pigment production gene (*crtM*) by binding to promoters of the corresponding factor genes. To test this hypothesis, promoters of the three gene operons were amplified by PCR and used for gel shift assay. As expected, the mobility of the three promoters became more hindered with increasing concentrations of the truncated RpvA, GB1-VpmR-S1, and formed obvious bands with lower electrophoretic mobility. Meanwhile, VpmR-S1 did not bind to *yacG* coding sequence (Figure 13 B).

**Figure 13.**
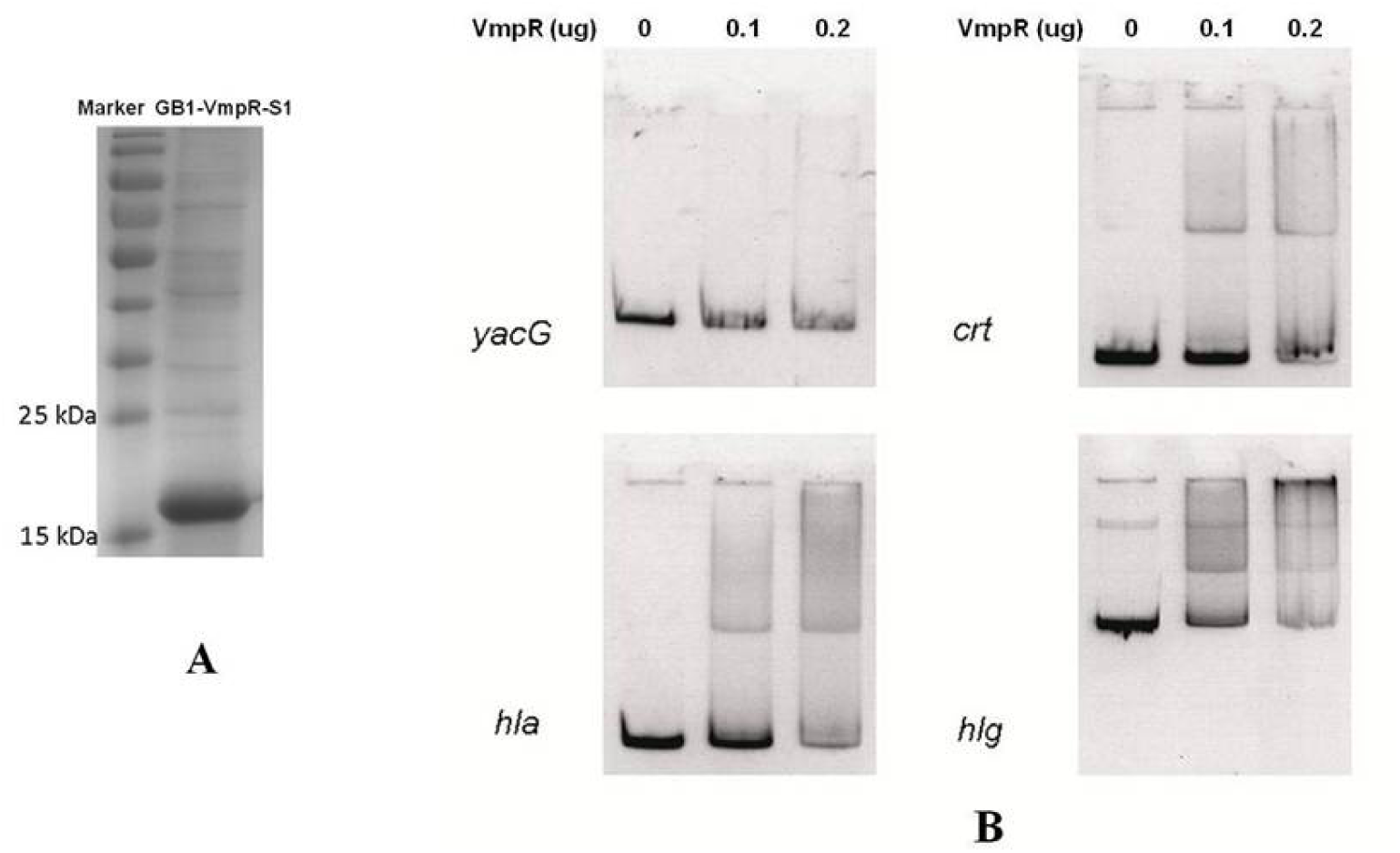
Purification of GB1-VpmR-S1 and gel shift assay. A. Purification of GB1-VpmR-S1. The molecular weight of about 18 kDa. B. Gel shift assay of VpmR. Promoter regions of indicated genes promoters were prepared by PCR. Each lane contained 10 ng DNA segments, which was co-cultured with 0, 0.1 and 0.2 μg GB1-VpmR-S1 respectively at 30 °C for 25 min before loading into the polyacrylamide gel.

## Discussion

Despite some progress is made about the mechanisms of persistence in *S. aureus* in recent years, the exact mechanisms by which *S. aureus* forms persisters as the culture transitions from log phase to stationary phase remains unclear. In this study, we identified a novel transcription factor, RpvA whose mutation caused a global defect in persistence to various antibiotics and stresses with attenuation of virulence. RpvA is a novel LysR family transcriptional regulator and its function was previously unknown. Transcriptional regulators can impact persister metabolism and cell survival (62, 63). LysR-type transcriptional regulators (LTTRs) are widely present in prokaryotes and can regulate a diverse set of genes, including those involved in virulence, metabolism, cell death, quorum sensing and motility in *S. aureus* (62, 64). Our study showed that RpvA is a novel transcription factor and an important global regulator for cellular metabolism, which allows *S. aureus* to form persisters and maintain stress resistance through regulating global metabolism including carbohydrate and amino acid metabolism, energy metabolism and metabolism of cofactors and vitamins, and enhance pathogenesis by increasing virulence factor expression in *S. aureus*.

Faced with environmental challenges such as antibiotics, stresses and nutrient deficiency, bacteria can adjust the cellular metabolism in a timely manner to form dormant persister cells, which can tolerate high stress levels and bactericidal effect of antibiotics (5, 6, 65). Previous studies have shown that many metabolic pathways such as carbohydrate metabolism, lipid, nucleotide and amino acid metabolism are involved in the formation of persisters (66, 67). Based on the transcriptomics and metabolomics analyses, we systematically analyzed the pathways by which RpvA could regulate the metabolism of *S. aureus* (Figure 14). Our work confirmed that the *rpvA* mutant had reduced persister formation in the early stationary phase (before 10 hours culturing). The major findings of RNA-seq revealed that at 6-hour culture stage, there were 73 genes upregulated but only 38 genes down-regulated in *rpvA* mutant compared with *S. aureus* Newman wild type (Table 4). Most upregulated genes (*tpi*, *gapR*, *gapA*, *pgm*, *eno*, *acs*, *glmS*, *glpD*, *narJ*, *narG*, *narH*, *alsS*) belonged to pathways including global metabolism, carbohydrate metabolism, energy metabolism, lipid metabolism, nucleotide metabolism, amino acid metabolism, metabolism of cofactors and vitamins and others, genetic information processing, environmental information processing (Figure 6 and 7), while 38 down-regulated genes are involved in virulence (*hla*, *hlgB*, *crtI*, *crtN*, *crtM*, *set1nm*) and metabolism (*pflB*, *hutG*, *mtlD*, *mtlA*, etc.). This indicates that RpvA mainly represses the bacterial cellular metabolism before the early stationary phase during the culture process, thereby more likely to be involved in formation of persisters rather than the persister survival. Similar phenomena and patterns were also confirmed by subsequent metabolomics. Analysis of all 76 differential metabolites between the *rpvA* mutant and wild type strain at the same time as RNA-seq revealed that 71 metabolites were decreased in the *rpvA* mutant, and only 5 species were increased, confirming that the *rpvA* mutation caused the metabolism of the bacteria to be highly active, which is reminiscent of the PhoU mutant which is a global regulator involved in persister formation in *E. coli* (Li and Zhang, 2007). In contrast, RpvA in the wild type bacteria presumably serves to inhibit bacterial metabolism, resulting in slow metabolism and accumulation of metabolites that would facilitate formation of persisters (Figure 6, 7, 9, and 10. Table 4 and 5).

**Figure 14.**
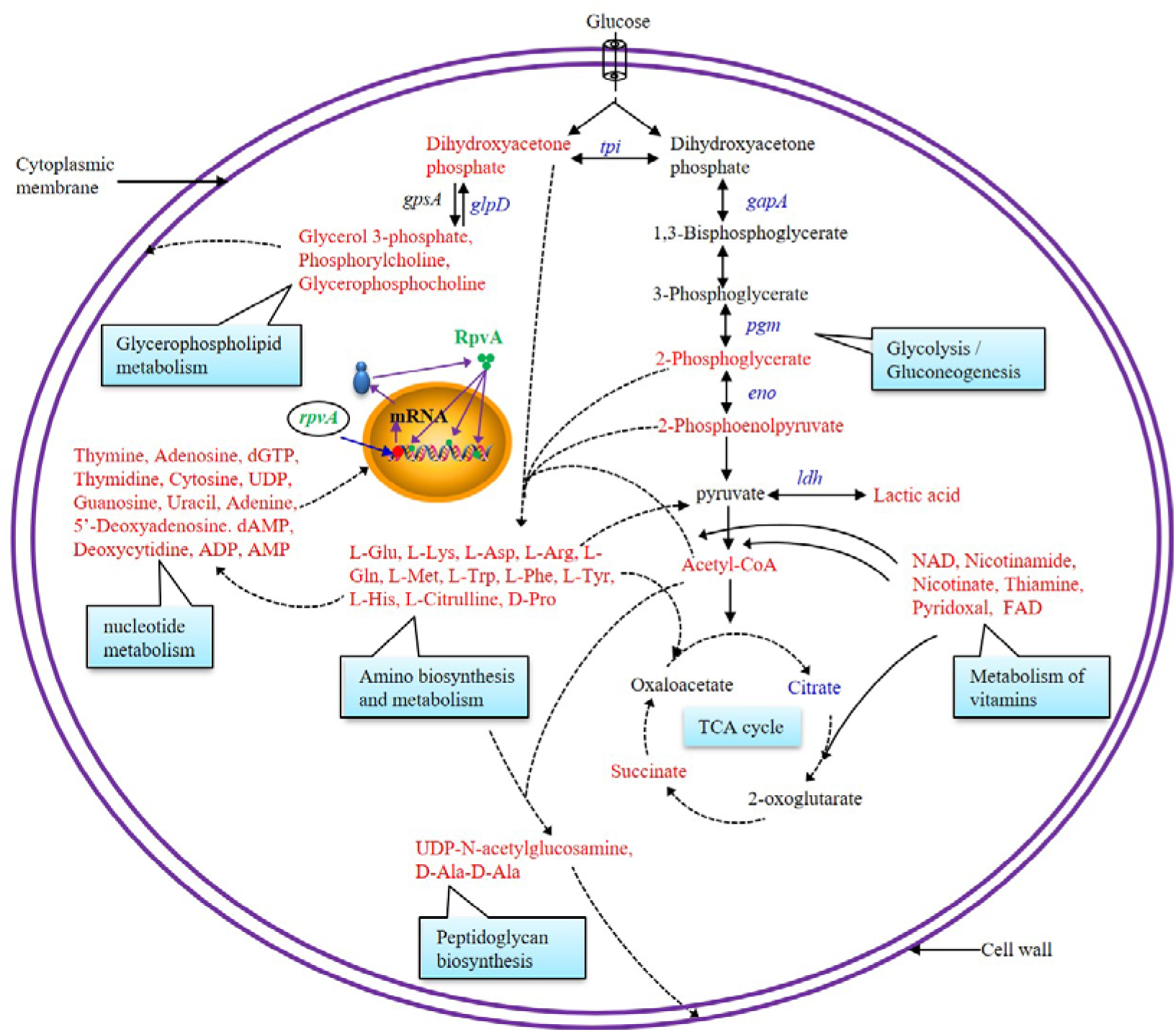
Main metabolism pathways that RpvA regulates in *S. aureus* Newman strain. RpvA is involved in the regulation of *S. aureus* glycolysis/gluconeogenesis, TCA cycle, amino acid biosynthesis and metabolism, glycerophospholipid metabolism, nucleotide metabolism, vitamin metabolism, peptidoglycan synthesis. Red substances represent significantly decreased metabolites and blue substances represent significantly upregulated genes and elevated metabolite in *rpvA* mutant compared with wild type.

Carbohydrate metabolism is the main pathway for bacteria to obtain energy, and decreased ATP production can promote persister formation (29–31). Most common metabolic pathway for bacteria utilizes glucose catabolism via glycolysis or Embden-Meyerhof-Parnas (EMP). Glyceraldehyde 3-phosphate dehydrogenase (GapA), phosphoglycerate kinase (Pgk), triosephosphate isomerase (Tpi), phosphoglyceromutase (Pgm), and enolase (Eno) which are encoded by *gapA*, *pgk*, *tpi*, *pgm*, and *eno* respectively are very important enzyme in glycolysis and gluconeogenesis pathways 68). These genes are regulated by a same operon named glycolytic operon regulator (*gapR*) in Newman strain DNA (69). Our work discovered that the *gapA*, *pgk*, *tpi*, *pgm*, *eno* and their operon *gapR* expression levels of *S. aureus* were significantly increased after mutation of *rpvA*. Correspondingly, the substrates catalyzed by these enzymes or the related products (Phosphoenolpyruvate, Dihydroxyacetone phosphate, DL-lactate, DL-2-Phosphoglycerate) were significant decreased in metabolite analysis. These results fully demonstrate that *rpvA* mutation significantly activates the glycolytic pathway, suggesting that RpvA may be a suppressor of glycolytic metabolism. Further research on the regulation of *gapR* and related glycolytic genes by RpvA is needed in the future. Acetyl coenzyme A and succinate are important ingredients of the tricarboxylic acid (TCA) cycle, and their significant decrease in *rpvA* mutant indicates that RpvA has an inhibitory effect on TCA. Previous studies have confirmed that inactivation of TCA cycle enhances *S. aureus* persister formation in stationary phase (39). Inhibition of glycolysis and TCA cycle reduces bacterial ATP production and promotes bacterial transformation to persister (29–31).

Our transcriptional profiling and metabolomic analysis found that RpvA is also involved in amino acid synthesis and metabolism, lipid, nucleotides, and vitamin metabolism, and peptidoglycan synthesis (Figure 6, 7, 9, and 10. Table 4 and 5). It is interesting that *glpD* whose product Glycerol-3-phosphate (G3P) dehydrogenase catalyzes G3P transformation in the *rpvA* mutant is significantly increased, accompanied by a decrease in G3P and dihydroxyacetone phosphate (DHAP) compared with the wild type (Figure 14). Previous studies have confirmed that null mutations in *glpD* increased the persister formation in *P. aeruginosa* and *E. coli* (70, 71). GlpD and G3P synthase (GpsA) encoded by *gpsA* function at the juncture between respiration and glycolysis, and catalyze the interconversion of G3P and DHAP. Decreased expression of *glpD* can cause accumulation of G3P, which in turn inhibits GpsA by a negative feedback mechanism, and eventually cause accumulation of DHAP. Elevated DHAP levels could produce methylglyoxal which could be responsible for the high-persistence frequency observed (70). GlpD is also an important enzyme involved in the glycerol metabolism pathway. The results are also consistent with our previous finding that GlpF involved in glycerol metabolism is associated with persister formation in *S. aureus* (37). Other phospholipid metabolism-related metabolites including phosphorylcholine and glycerophosphocholine which are linked with bacterial membrane synthesis were also decreased in the *rpvA* mutant compared with wild type. Integrity and permeability of cytoplasmic membrane affect persister sensitivity to antimicrobials (72) and could explain the increased susceptibility of the RpvA mutant to antibiotics and stresses.

A significant decrease in the content of various amino acids and nucleotides were identified in the *rpvA* mutant compared with wild type, demonstrating that RpvA is involved in the biosynthesis and metabolism of amino acids and nucleotides (Figure 6, 7, 9, and 10. Table 4 and 5). The biosynthesis and metabolism of many amino acids and nucleotides which participate in persister formation had been confirmed in previous studies. Carbamoyl phosphate synthetase (CPSase) is a metabolic enzyme involved in the synthesis of *P. aeruginosa* pyrimidine and arginine, and mutation of *carB* encoding the large subunit of CPSase resulted in a 2500-fold reduction in survival after antibiotic treatment, while addition of uracil could abolish this phenotype. This indicates that the biosynthesis of arginine and uracil is closely related to the formation of persister (73). Faced with reduced sugar availability in the environment, *S. aureus* could catabolize multiple amino acids as a secondary carbon source to generate pyruvate (alanine, serine, glycine, threonine, cysteine), 2-oxoglutarate (glutamate, glutamine, histidine, arginine, proline), and oxaloacetate (aspartate, asparagine) (74). Glutamate dehydrogenase (GudB) can convert glutamate into 2-oxoglutarate which can fuel the TCA cycle under glucose-depleted conditions (74). Knock out of *gudB* can decrease ATP levels in late exponential phase *S. aureus* and enhance tolerance to antibiotics significantly (24). A significant decrease of L-tryptophan, L-arginine, L-glutamate, L-glutamine, L-histidine, L-aspartate, and D-proline identified in the *rpvA* mutant could underlie the defect in persistence as seen in this study.

Peptidoglycan synthesis is related to the formation of persister (Moyed HS, Bertrand KP. J Bacteriol. 1983 Aug;155(2):768-75), and it has been shown that inhibition of peptidoglycan synthesis contributes to persister formation (75). Our study showed that UDP-N-acetylglucosamine and D-Ala-D-Ala, which are involved in peptidoglycan synthesis, were significantly decreased in the *rpvA* mutant, which may not be conducive to the formation of persisters. It is well known that staphyloxanthin function as antioxidants which can protect *S. aureus* against oxidative stress (76). Mutation of *rpvA* leads to down-regulation of genes involved in staphyloxanthin biosynthesis including dehydrosqualene synthase (*crt*M), desaturase (*crt*N) and phytoene dehydrogenase (*crt*I), which could cause the mutant to be more sensitive to the peroxide stress as seen in this study.

Cytotoxins, enterotoxin, exfoliative toxin and coagulase are major virulence factors which mediate *S. aureus* infections. Our research showed that RpvA acts as a positive regulator for virulence factors in *S. aureus* (Figure 15). These can explain why the Newman wild type survived better in macrophages, had lower LD50 and higher pathogenicity in the murine skin abscess model than the *rpvA* mutant (Figure 4 and 5). Consistent with the above finding, qPCR study showed that the major virulence genes (*hla*, *hlgA*, *hlgB*, *hlgC*, *lukS*, *lukF*, *lukD*, *sea*, *coa*, and *eta*) of Newman wild type were significantly upregulated compared with the *rpvA* mutant (Figure 12). Gel shift assay confirmed RpvA fragment binds to promoters of virulence genes *crtM*, *hla* and *hlgB* directly (Figure 13). *hla* encodes α-hemolysin which can bind to the cell surface and assemble into a homoheptamer and form a pore which allows transport of K^+^ and Ca^2+^ ions, leading to necrotic death of the target cell (42). α-hemolysin is susceptible to human erythrocytes, lymphocytes, and monocytes (42, 77), which can cause hemolysin and damage human’s immune function. *hlg*A, *hlg*B, and *hlg*C encode γ-hemolysin which is composed with a slow component (HlgA or HlgC) and a fast component (HlgB). *luk*S and *luk*F encode PVL which made of LukS-PV and LukF-PV (78). γ-hemolysin and PVL are highly toxic to human neutrophils, monocytes and macrophages which can also damage and inhibit the immune function (79, 80). Coagulase can convert fibrinogen to insoluble fibrin and cause the formation of a fibrin layer around a staphylococcal abscess, thus protecting the bacteria from phagocytosis. Exfoliative toxins are serine protease that split the intercellular bridge and cause staphylococcal scalded skin syndrome. Staphylococcal enterotoxins are heat-stable toxins associated with food poisoning. Enterotoxins are superantigens which can induce nonspecific activation of T cells and massive cytokine release (46). The higher expression of genes for coagulase, exfoliative toxins, and enterotoxins in the wild type Newman strain compared with the *rpvA* mutant can explain the higher virulence in the wild type Newman strain but attenuated virulence in the *rpvA* mutant. In addition, we found that RpvA can enhance carotenoid staphyloxanthin production which help the microbe copy with reactive oxygen species (ROS) generated by neutrophils and macrophages, thereby enhancing the virulence and fitness of the cells (Figure 15) (76, 81).

**Figure 15.**
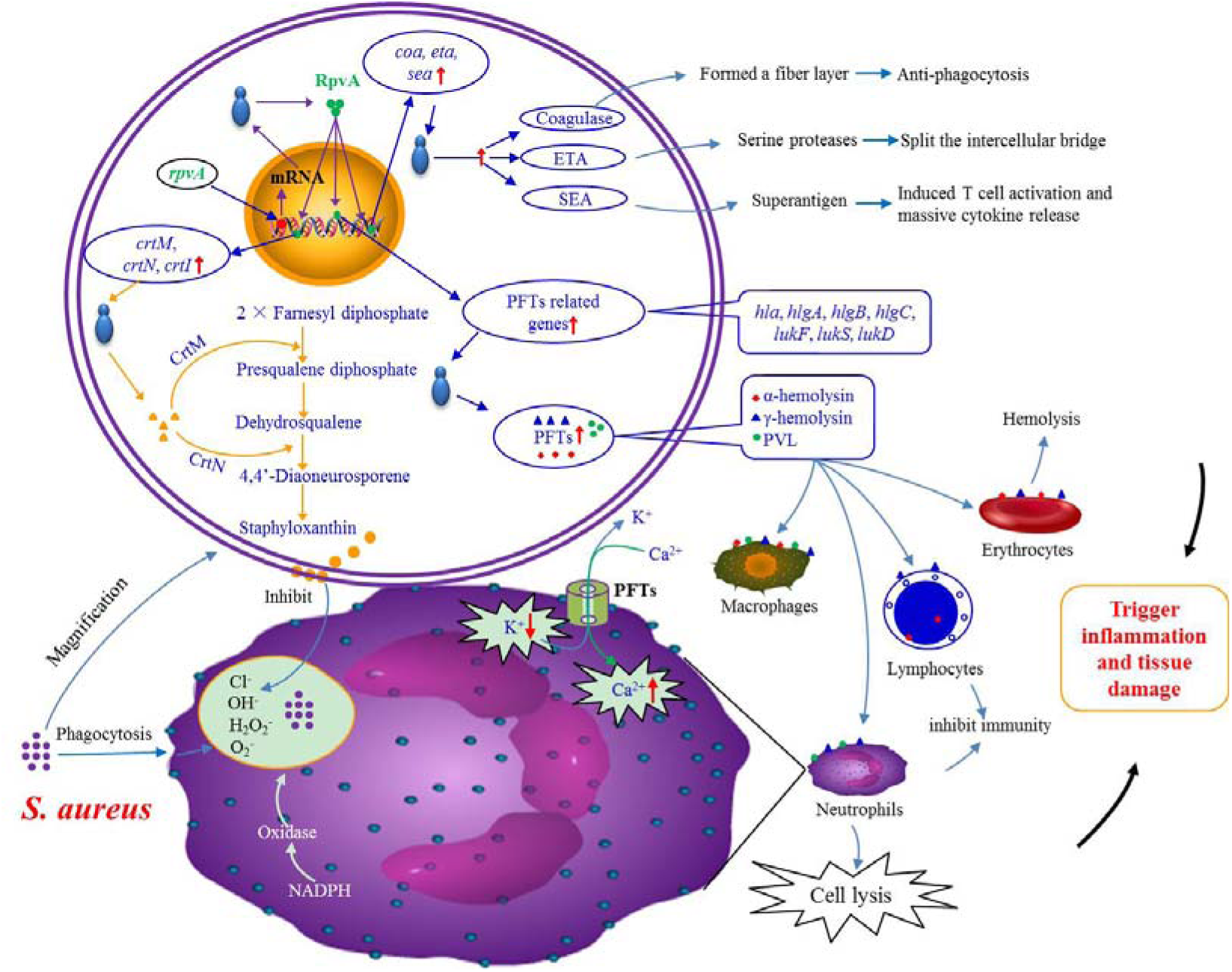
Mechanisms by which RpvA regulates virulence of *S. aureus* Newman strain. Red upward arrows indicate increase or upregulated, and red downward arrow indicates decrease or downregulated. RpvA can upregulate toxin related gene (*hla*, *hlgA*, *hlgB*, *hlgC*, *lukS*, *lukF*, *lukD*, *coa*, *sea*, and *eta*) expression. The products including α-hemolysin, γ-hemolysin, and PVL can damage the leukocytes, neutrophils, macrophages and erythrocytes, which can destroy erythrocyte and host immunity. Coagulase can protect the bacteria from phagocytosis. Exfoliative toxins (ETA) can split the intercellular bridge. Staphylococcal enterotoxins (SEA) can induce nonspecific activation of T cells and massive cytokine release. RpvA can enhance *crtM*, *crtN*, and *crtI* expression which can affect carotenoid staphyloxanthin synthesis, and helps the microbe to promote resistance to ROS generated by neutrophils and macrophages and host immune system. All these interactions trigger inflammation and cause tissue damage.

In summary, we identified a novel LysR-type transcription factor RpvA that is a global regulator and plays an important role in persister formation and virulence regulation in *S, aureus*. RpvA is closely related to *S. aureus* exponential and early stationary phase persisters formation, stress response and enhancing virulence through regulation of global metabolism including carbohydrate metabolism, amino acid metabolism, energy metabolism and metabolism of cofactors. Further studies are needed to determine how the different mechanisms cooperate to mediate persister formation, stress response and virulence in response to environmental cues. Because RpvA is involved in persister formation, stress response and enhancing virulence, it should serve as an attractive target for developing persister drugs and vaccines for more effective control of *S. aureus* infections.

## Acknowledgments

Jian Han was sponsored by China Scholarship Council. This research was supported by National Natural Science Foundation of China (84150043, 81571952).

